# Foveated metamers of the early visual system

**DOI:** 10.1101/2023.05.18.541306

**Authors:** William F. Broderick, Gizem Rufo, Jonathan Winawer, Eero P. Simoncelli

## Abstract

The ability of humans to discriminate and identify spatial patterns varies across the visual field, and is generally worse in the periphery than in the fovea. This decline in performance is revealed in many kinds of tasks, from detection to recognition. A parsimonious hypothesis is that the representation of any visual feature is blurred (spatially averaged) by an amount that differs for each feature, but that in all cases increases with eccentricity. Here, we examine models for two such features: local luminance and spectral energy. Each model averages the corresponding feature in pooling windows whose diameters scale linearly with eccentricity. We performed perceptual experiments with synthetic stimuli to determine the largest window scaling for which human and model discrimination abilities match (the “critical” scaling). We used much larger stimuli than those of previous studies, subtending 53.6 by 42.2 degrees of visual angle. We found that the critical scaling for the luminance model was approximately one-fourth that of the energy model and, consistent with earlier studies, that the estimated critical scaling value was smaller when discriminating a synthesized stimulus from a natural image than when discriminating two synthesized stimuli. Moreover, we found that initializing the generation of the synthesized images with natural images reduced the critical scaling value when discriminating two synthesized stimuli, but not when discriminating a synthesized from a natural image stimulus. Together, the results show that critical scaling is strongly affected by the image statistic (pooled luminance vs. spectral energy), the comparison type (synthesized vs. synthesized or synthesized vs. natural), and the initialization image for synthesis (white noise vs. natural image). We offer a coherent explanation for these results in terms of alignments and misalignments of the models with human perceptual representations.

## Introduction

Vision science is often concerned with what things look like (“appearance”), but a long and fruitful thread of research has investigated what humans cannot see, that is, the information they are insensitive to. Perceptual metamers — images that are physically distinct but perceptually indistinguishable — provide a classic example of such research, used to elucidate loss of visual information. Cohen and Kappauf (1985) identified this concept in the writings of Isaac Newton, who noted that the color percept elicited by a single wavelength of light could also be created by mixing multiple wavelengths. Cohen and Kappauf traced the perceptual meaning of the word “metamer” to a 1919 chapter by Wilhelm Ostwald, the first Nobel laureate in chemistry. Color metamers were instrumental in the development of the Young-Helmholtz theory of trichromacy (Helmholtz, 1852). Specifically, the experimental observation of metamers clarified human sensitivity to light wavelengths, and led to the hypothesis that the human visual system projects the infinite-dimensional physical signal to three dimensions. It took more than a century before the physiological basis for this — the three classes of cone photoreceptor — was revealed experimentally (Schnapf et al., 1987). This projection into a three-dimensional space enables one to predict which lights will look identical. It makes no prediction of how discriminable lights will be when they are not metameric. Predicting perceptual differences within the color-matching space remains an ongoing effort (Brainard, 2022).

The visual system also discards a great deal of spatial detail, more so in portions of the visual field farthest from the center of gaze. Specifically, the reduction of visual capabilities with increasing eccentricity has been demonstrated for acuity (Frisen and Glansholm, 1975) and recognition (Pelli and Tillman, 2008), and is reflected in the physiology: fewer cortical resources are dedicated to the periphery (Schwartz, 1977) and receptive fields in all stages of the visual hierarchy grow with eccentricity (e.g., Daniel and Whitteridge (1961); Dacey and Petersen (1992); Gattass et al. (1981, 1988); Maunsell and Newsome (1987); Wandell and Winawer (2015)). This decrease in acuity has been demonstrated by either scaling the size of features, as in the Anstis eye chart (Anstis, 1974), or by progressively blurring the image, as in Anstis (1998); Thibos (2020). More generally, one can explain this decreasing sensitivity to spatial information with “pooling models”, which compute local averages of image features in windows that grow larger with eccentricity (Balas et al., 2009; Freeman and Simoncelli, 2011; Keshvari and Rosenholtz, 2016). These models assume that peripheral representations are qualitatively similar to those of the fovea: the same local computations are performed over larger regions, leading to greater dimensionality reduction in the periphery.

Such pooling at increasingly larger scales is ubiquitous throughout the many stages of visual processing. One hypothesis is that each stage performs essentially the same canonical computation, “extract features and pool”, differing based on what features are extracted and the spatial extent of the pooling. Here, we test models based on two kinds of image features — one model that averages local luminance (luminance model) and one that averages both local spectral energy and luminance (energy model). We hypothesize that the more complex feature, spectral energy, is pooled over a larger spatial extent than luminance. We test the spatial pooling extent of these image features using a metamer paradigm. Specifically, we generate image pairs in which one or both images have been manipulated such that the two are model metamers (images that are physically distinct but with identical model representations). The pair of model metamers are also perceptual metamers if the human visual system is insensitive to the differences between them, as schematized in figure 1 (see Watson et al. (1986) for an analogous presentation with respect to sensitivity to spatial and temporal frequency) ^1^. By comparing model and human perceptual metamers, we investigate how well the models’ sensitivities (and insensitivities) align with those of the human visual system. Note that throughout much of this paper, we refer to “visual stimuli” to emphasize that our results and interpretation depend on the viewing conditions (distance from viewer, relative location of the fovea, and linear relationship between pixel values and output luminance).

**Figure 1.**
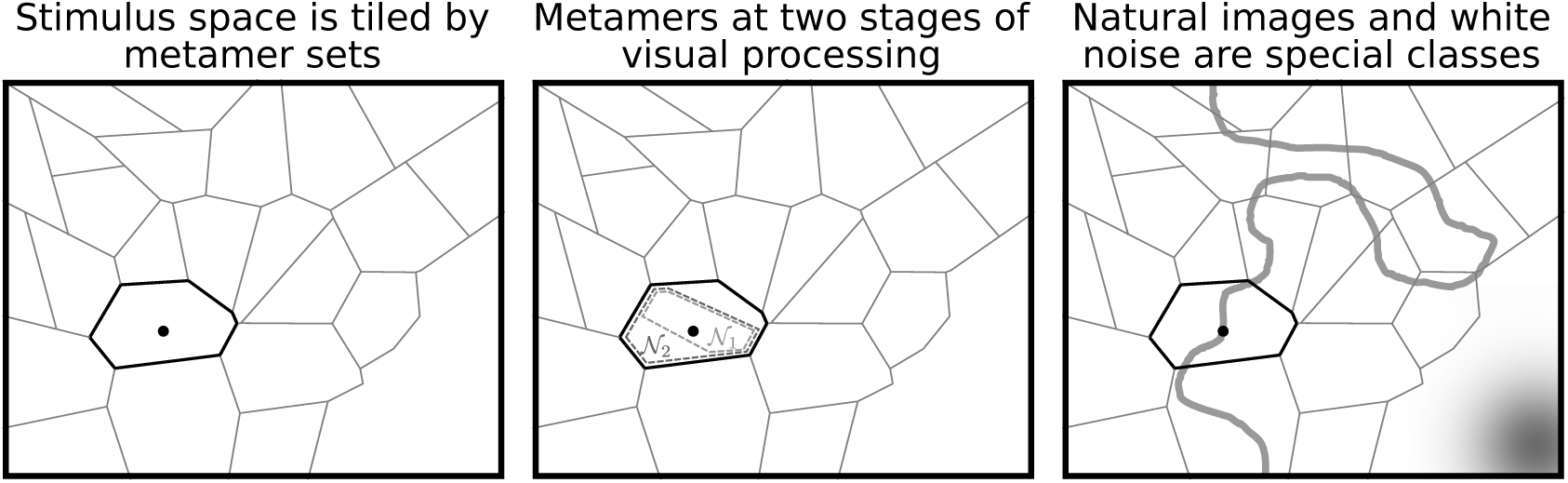
Idealized conceptual schematic of perceptual metamers. Each panel contains a two-dimensional depiction of the set of all visual images: every image corresponds to a point in this space and every point in this space represents an image. In this and similar figures, each tile represents the set of images which project to the identical point in a perceptual space and hence are metameric, analogous to different lights projecting to the same tri-stimulus coordinates in a color matching space. Any pair of points in different tiles are assumed to project to different locations in a perceptual space; their discriminability depends on the internal signal-to-noise level, analogous to neighboring points in a color matching space. Left: example stimulus (black point), and surrounding metameric stimuli (region enclosed by black polygon). Center: In a hierarchical visual system, in which each stage transforms the signals of the previous stage and discards additional information, every pair of stimuli that are metamers for an early visual area N_1_ are also metamers for the later visual area *N*_2_. Thus, metamers of earlier stages are nested within metamers of later stages. The converse does not necessarily hold: there may be stimuli that *N*_1_ can distinguish but that *N*_2_ cannot. Right: Two stimulus families used for initialization of our metamer synthesis algorithm: white noise (distribution represented as a grayscale intensity map in the lower right corner) and natural images (distribution represented by the curved gray manifold). Typical white noise samples fall within a single perceptual metamer class (humans are unable to distinguish them). Natural images, on the other hand, span many tiles in the perceptual space, and are generally distinguishable from each other.

This procedure rests on the assumption that the visual system processes information hierarchically, in a sequence of stages, and information discarded in early stages cannot be recovered by later stages (the “data-processing inequality”) ^2^. For example, metameric color stimuli produce identical cone responses and thus cannot be distinguished by any additional downstream neural processing. Similarly, if two visual stimuli generate identical responses in all neurons at a subsequent stage of processing (e.g., the retinal ganglion cells), the stimuli will appear identical, even if their cone responses differ. This is schematized in the central panel of figure 1: two stimuli are perceptual metamers if they are indistinguishable in an early visual area, such as *N*_1_ or *N*_2_.

Combining these pooling models with the metamer paradigm allows us to ask over which spatial scales are humans sensitive to changes in a given statistic. The model metamers are defined as the set of visual stimuli that match a target visual stimulus in the specified image statistic averaged at the spatial scale defined by the model’s scaling parameter, with (ideally) all other aspects of image content as random as possible. If humans are sensitive to the image statistic at a scale smaller than the one used to synthesize the model metamer, then they may be able to tell the difference between the model metamer and the target image, or to pairs of model metamers that are physically distinct. If humans are sensitive to the image statistic only at the same or larger spatial scale, then they will be unable to distinguish between the set of stimuli, and thus the stimuli are also perceptual metamers.

A number of authors have investigated perceptual metamers using pooling models, using a particular family of “texture statistics” capturing joint responses of oriented filter responses and their nonlinear combinations (e.g., Freeman and Simoncelli (2011); Keshvari and Rosenholtz (2016); Wallis et al. (2019); Deza et al. (2019); Brown et al. (2023)). In this study, we instead investigate models based on local pooling of either luminance or spectral energy. These are arguably the two most fundamental attributes of visual stimuli, and they are commonly used to characterize inputs (visual stimuli) and outputs (neural responses and behavior, as in neurometric and psychometric functions of luminance and contrast). Moreover, these two measures correspond to the mean (luminance) and the variance within frequency channels (spectral energy), the two most common choices for statistical summary of central tendency and dispersion. While humans are sensitive to both measures, neither is intended as a model of specific neural responses in the human visual system. The use of these relatively simple statistics also facilitates experimental design and interpretation. For example: how should one best construct the set of model metamer stimuli to use in the experiment? How should one reconcile differing results that arise from comparing two synthesized stimuli or one synthesized stimulus with the original image stimulus?

The importance of these questions has been highlighted in two previous studies: Wallis et al. (2019) found that performance was poorer when participants compared two synthesized stimuli than when they compared a synthesized stimulus to the original image, and Brown et al. (2023) observed performance differences dependent on base image characteristics. Here, we examine both of these factors, as well as the influence of the distribution of “seed” images used to initialize the stochastic metamer synthesis algorithm. In general, the model metamers synthesized for use in any experiment represent only a small subset of all such metamers, and the procedure by which those samples are generated can have a strong influence on the outcome.

In this study, we synthesized metamers for two different models at multiple spatial scales and measured their perceptual discriminability. For a set of 20 natural images, we used a stochastic gradient descent method to generate metamers for both models, and measured discrimination capabilities of human observers when comparing these with their corresponding original images, as well as with each other. Finally, we compared our results with those of previous studies that used a texture model whose statistics were pooled in eccentricity-scaled regions (Freeman and Simoncelli, 2011; Wallis et al., 2019), and found that they were consistent.

## Results

### Foveated pooling models

We constructed foveated models of human perception that capture sensitivity to local luminance and spectral energy (see figure 2). Both models are “pooling models” (Balas et al., 2009; Freeman and Simoncelli, 2011; Keshvari and Rosenholtz, 2016; Wallis et al., 2019), which compute statistics as weighted averages within overlapping local windows. A specific pooling model is characterized by both the quantities that are pooled and the shapes/sizes of the pooling windows. These regions are analogous to neural receptive fields, whose sizes grow proportionally with distance from the fovea, as documented in monkey physiology and human fMRI (e.g., Gattass et al. (1981); Wandell and Winawer (2015)).

**Figure 2.**
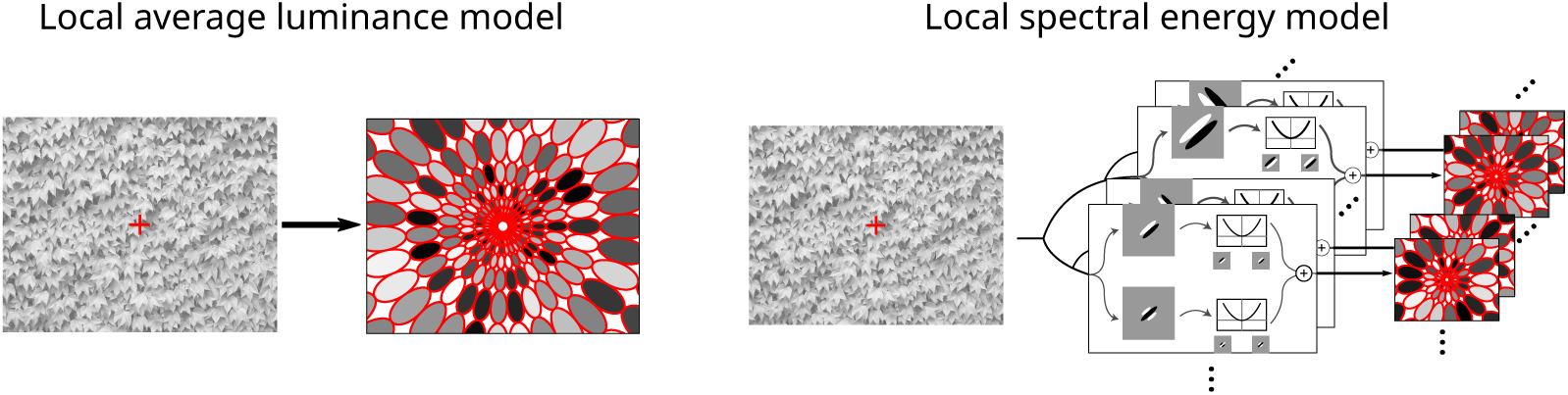
Two pooling models. Both models compute local image statistics within a Gaussian-weighted window that is separable in a log-polar coordinate system, such that radial extent is approximately twice the angular extent (red contours indicate half-maximum levels). Windows are uniformly spaced in a log-polar grid, with centers separated by one standard deviation. A single scaling factor governs the size of all pooling windows. The luminance model (left) computes average luminance. The spectral energy model (right) computes average spectral energy at 4 orientations and 6 scales, as well as luminance, for a total of 25 statistics per window. Spectral energy is computed using the complex steerable pyramid constructed in the Fourier domain (Simoncelli and Freeman, 1995), squaring and summing across the real and imaginary components. Full resolution version of this figure can be found on the OSF.

As in previous studies (Freeman and Simoncelli, 2011; Keshvari and Rosenholtz, 2016; Wallis et al., 2019), we used arrays of smooth overlapping windows that are separable and of constant size when expressed in polar angle and log-eccentricity (consistent with the approximate log-polar geometry of visual cortical maps (Schwartz, 1977)), as illustrated in figure 2. Like these previous studies, our windows were radially-elongated (the radial extent is roughly twice the angular extent), but unlike the previous studies, we used Gaussian profiles separated by one standard deviation, yielding a smoother representation and minimal ringing and blocking artifacts in the synthesized stimuli (see appendix 3 for a comparison of the two window profiles). The log-polar mapping ensures that the size of the pooling windows, in both radial and angular directions, is proportional to their distance from the fovea, and this proportionality is controlled by a single scaling parameter, *s*. Increasing this value increases the size of all pooling regions, resulting in a larger and more diverse set of metameric stimuli, in a nested manner (figure 3).

**Figure 3.**
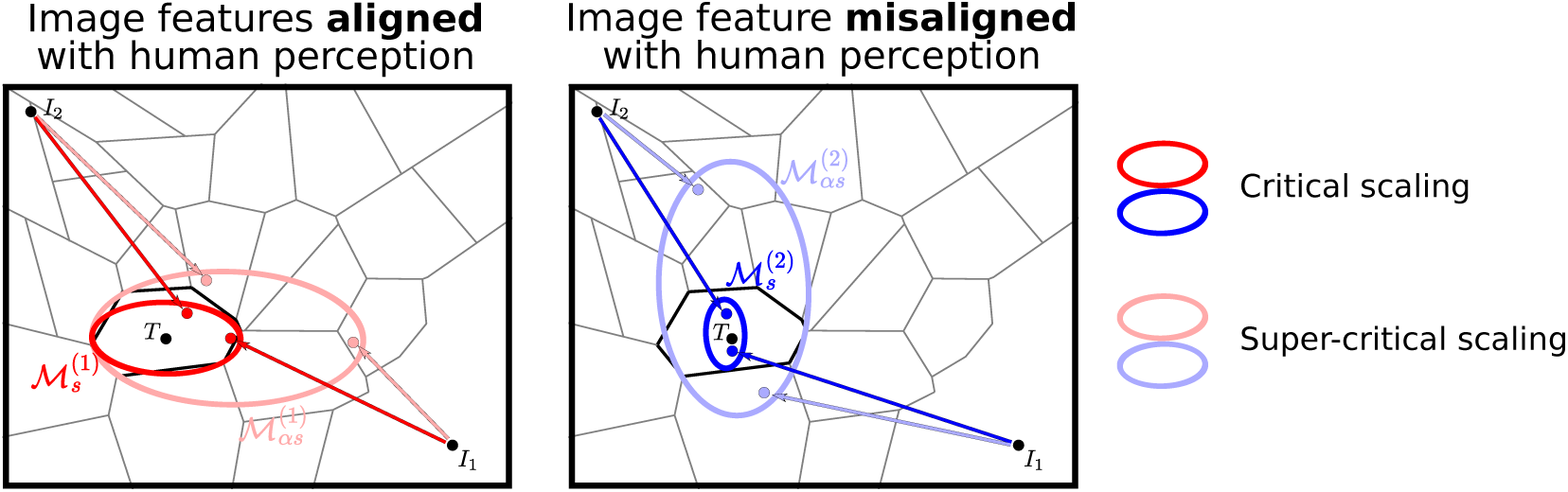
Metamers in human perception and pooling models. Panels depict a two-dimensional stimulus space, with polygonal regions indicating groups of perceptually indistinguishable stimuli (human metamers, see figure 1). Pooling models n^(^*^i^*^)^ determine the statistics that are pooled, with pooling extent controlled by scaling parameter, *s*. Ellipses indicate *model metamers*: sets of stimuli that are physically different but whose model outputs are identical to those of the target (original) image *T*. We generate model metamer samples using an optimization procedure: starting from initial image *I_k_* we adjust the pixel values until their pooled statistics match those of the target. The shapes and sizes of metameric regions (ellipses) depend on the model (*i*), the scaling parameter (*s*), the statistics of the target image *M^(i)^_s_(T)*, as well as the initial images (*I_k_*) and the synthesis algorithm. For a given set of statistics, increasing the scaling value by factor *α >* 1 increases the size of the metamer set, and any stimulus that is a metamer for *M^(i)^_s_* will also be a metamer for *M^(i)^_αs_* (i.e., larger desaturated ellipses contain the smaller saturated ellipses). *Critical scaling* is the largest *s* for which all model metamers are human metamers (smaller ellipses). Left: Model with parameters that produce approximate alignment with human metamers. Right: Model with different pooled statistics that yield metamers (blue ellipses) that are poorly aligned with human metamers: at critical scaling, there are human metamers that are not model metamers. This mismatch cannot be resolved by adjusting the scaling parameter: increasing *s* such that all human metamers are also model metamers (larger ellipse) will also yield model metamers that are not human metamers. See also Feather et al. (2023), figure 1.

In the current study, we examine two models that compute different statistics within their pooling windows. The luminance model pools pixel intensities, and thus, two luminance model metamers have the same average luminance within corresponding pooling windows. This model serves as a baseline for comparison to models that pool more image statistics, such as the energy model used here and the texture model used in Freeman and Simoncelli (2011); Wallis et al. (2019). Since the model’s responses are insensitive to the highest frequencies, luminance model metamers include blurred versions of the target image (in which high frequencies are discarded), but also variants of the target image in which high frequencies are randomized or even amplified. In general, synthesized luminance model metamers inherit the high frequency content of their initialization image, as can be seen in figure 4, middle row. While the high-scaling model metamer is clearly perceptually distinct from the target image (regardless of observer fixation location), the low-scaling stimulus is difficult to discriminate from the target when fixating at the center of the stimulus (i.e., when the human observer’s fovea is aligned with the model fovea).

**Figure 4.**
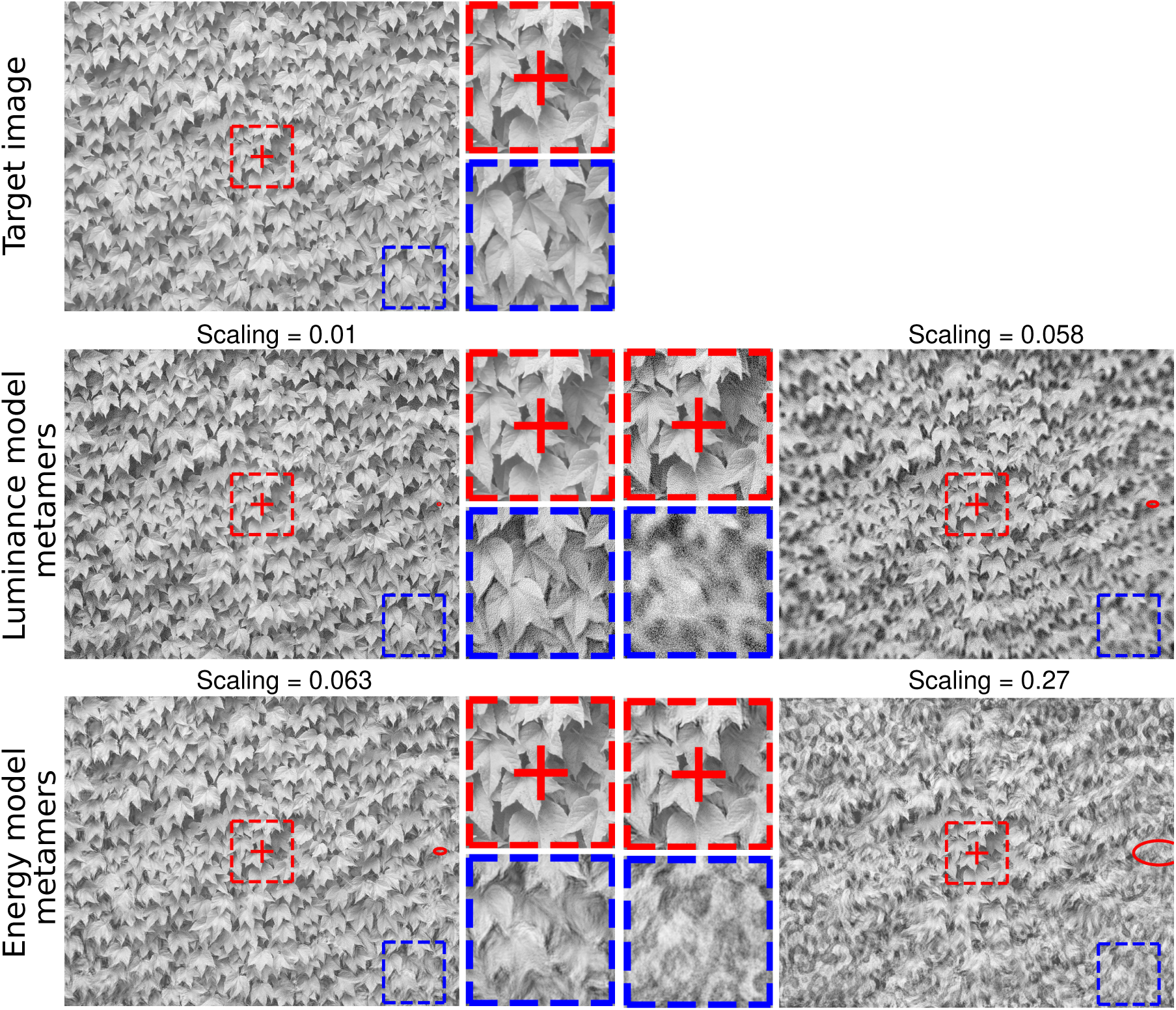
Example synthesized model metamers. Top: Target image. Middle: Luminance model metamers, computed for two different scaling values (values as indicated, red ellipses to right of fixation indicate pooling window contours at half-max at that eccentricity). The left image is computed with a small scaling value, and is a perceptual metamer for most subjects: when fixating at the cross in the center of the stimulus, the two stimuli appear perceptually identical to the target image. Note, however, that when fixating in the periphery (e.g., the blue box), one can clearly see that the stimulus differs from the target (see enlarged versions of the foveal and peripheral neighborhoods to right). The right image is computed with a larger scaling value, and is no longer a perceptual metamer (for any choice of observer fixation). Bottom: Energy model metamers. Again, the left image is computed with a small scaling value and is a perceptual metamer for most observers when fixated on the center cross. Peripheral content (e.g., blue box) contains more complex distortions, readily visible when viewed directly. The right image, computed with a large scaling value, differs in appearance from the target regardless of observer fixation. Full resolution version of this figure can be found on the OSF.

The spectral energy model pools the squared outputs of oriented bandpass filter responses at multiple scales and orientations, as well as the pixel intensities. As such, its statistics are a superset of the luminance model, and its metamers are a subset. The energies are computed using a complex steerable pyramid, which decomposes images into frequency channels selective for 6 different scales and 4 different orientations. Energy is computed by squaring and summing the real and imaginary responses (arising from even- and odd-symmetric filters) within each channel. These energies, along with the luminances, are then averaged within the spatial pooling windows. Thus, a pair of energy model metamers have the same average oriented energy and luminance within each of these windows. The bottom row of figure 4 shows energy model metamers for two different scaling values. The low scaling value for the energy model is approximately matched to the higher scaling value for the luminance model, while the higher scaling value is approximately that of the energy model from Freeman and Simoncelli (2011). The high-scaling model metamer is perceptually distinct from the target stimulus, and also perceptually distinct from the high-scaling luminance model metamer. The low-scaling model metamer, on the other hand, is difficult to distinguish from the original image (when fixating at the center), but is readily distinguished when one fixates peripherally.

The appearance of these two model metamers reflects both the measurements that are being matched and the seed images used to initialize synthesis. The luminance model matches average pixel intensity, but has no constraints on spatial frequency, and thus its metamers retain the high frequency content of the initial white-noise images. The energy model, on the other hand, matches the average contrast energy at all scales and orientations, but discards exact position information (which depends on phase structure). Hence, unlike the luminance model metamers, it reduces the high frequency power to match the typical content of natural images, and essentially scrambles the phase spectrum, leading to the cloud-like appearance of its metamers. For both models and all scales tested, the synthesized stimuli are physically very different from the target stimuli, as measured by mean squared error between the images (see appendix 5).

As can be seen in figure 4, both models can generate perceptual metamers. More generally, all pooling models can generate perceptual metamers if the scaling value is made sufficiently small (in the limit as scaling goes to zero, the model metamers must be identical to the target, in every pixel). For statistics that capture features relevant to human perception, metamers can be achieved with windows whose size is matched to the scale of human perceptual sensitivity. The maximal scaling at which synthetic stimuli are perceptual metamers is thus highly dependent on the choice of underlying statistics: in our examples, the energy model perceptual metamer (figure 4, bottom left) is generated with a scaling value about six times larger than that for the luminance model perceptual metamer (middle left), and about five times smaller than those reported in Freeman and Simoncelli (2011) using higher-order texture statistics. The goal of the present study is to use psychophysics to find the largest scaling value for which these two models generate perceptual metamers, known as the critical scaling.

### Psychophysical experiment

We synthesized model metamers matching 20 different natural images (the target images) collected from the authors’ personal collections, as well as from the UPenn Natural Image Database (Tkačik et al. (2011), extended dataset of urban images provided by David Brainard). The images were chosen to span a variety of natural image content types, including buildings, animals, and natural textures (figure 5). Model metamers were generated via gradient descent on the squared error between target and synthetic pooled statistics, and initialized with either an image of white noise or another image drawn from the set of target images.

**Figure 5.**
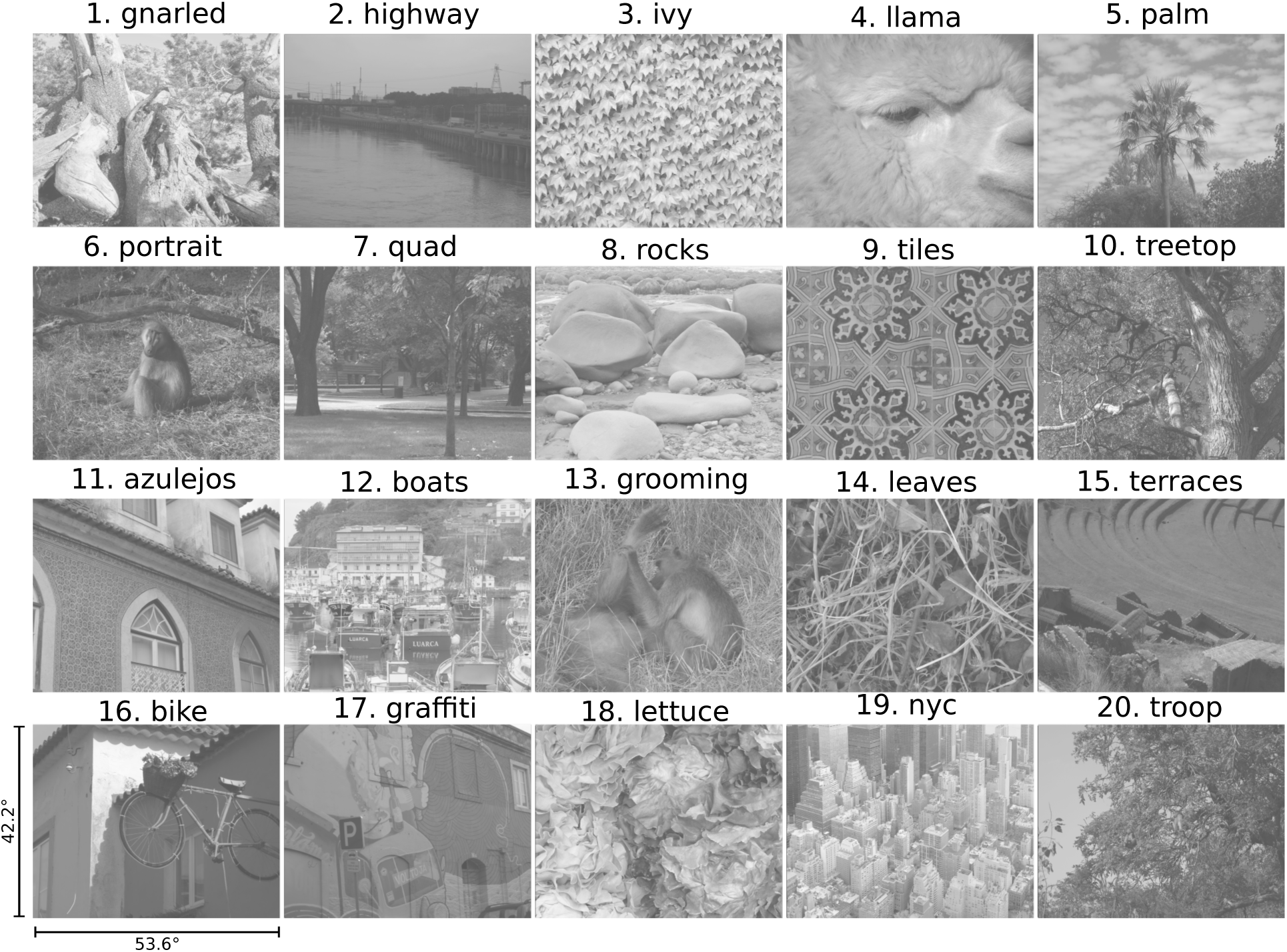
Target images used in the experiments. Images contain a variety of content, including textures, objects, and scenes. All are RAW camera images, with values proportional to luminance and quantized to 16 bits. Images were converted to grayscale, cropped to 2048 × 2600 pixels, displayed at 53.6 × 42.2 degrees, with intensity values rescaled to lie within the range of [0.05, 0.95] of the display intensities. All subjects saw target images 1–10, half saw 11–15, and half saw 16–20. A full resolution version of this figure can be found on the OSF.

In the experiments, observers discriminated two grayscale stimuli, of size 53.6 by 42.2 degrees, sequentially displayed. Each stimulus was separated into two halves by a superimposed vertical bar (mid gray, 2 deg wide, see figure 15). One side, selected at random on each trial, was identical in the two intervals, while the other differed (e.g., during the second interval, one half of the stimulus contains the target image, the other a synthesized model metamer). Each stimulus was presented for 200 msecs, separated by a 500 msec blank (mid gray) interval, and followed by a blank screen with text prompting the observer to report which side of the stimulus had changed.

There are several reasons to think that the subjects generally maintained good fixation (see Methods Apparatus section). However, we did not use eye tracking in the psychophysical experiment, and thus cannot guarantee that our observers maintained fixation for all trials.

### Critical scaling is four times smaller for the luminance than the energy model

We fit the behavioral data using the 2-parameter function introduced in Freeman and Simoncelli (2011), estimating the critical scaling (*s_c_*) and maximum *d*^f^ (*a*) parameters with a Markov Chain Monte Carlo procedure and a hierarchical, partial-pooling model similar to that used by Wallis et al. (2019). For a given model and comparison, performance increases monotonically with scaling, and is fit well by this particular psychometric function (figure 6A). The exception is the synthesized vs. synthesized comparison for the luminance model, for which performance remains poor at all scales (see next section). In the original vs. synthesized cases (for both models), performance is near chance for the smallest tested scaling values and exceeds 90% for the largest. The critical scaling values, as seen in figure 6B, are approximately 0.016 for the luminance model and 0.06 for the energy model. Thus, as expected, the critical scaling for the model depends on the statistic being pooled: humans are sensitive to changes in local luminance at a much smaller scale than local spectral energy.

**Figure 6.**
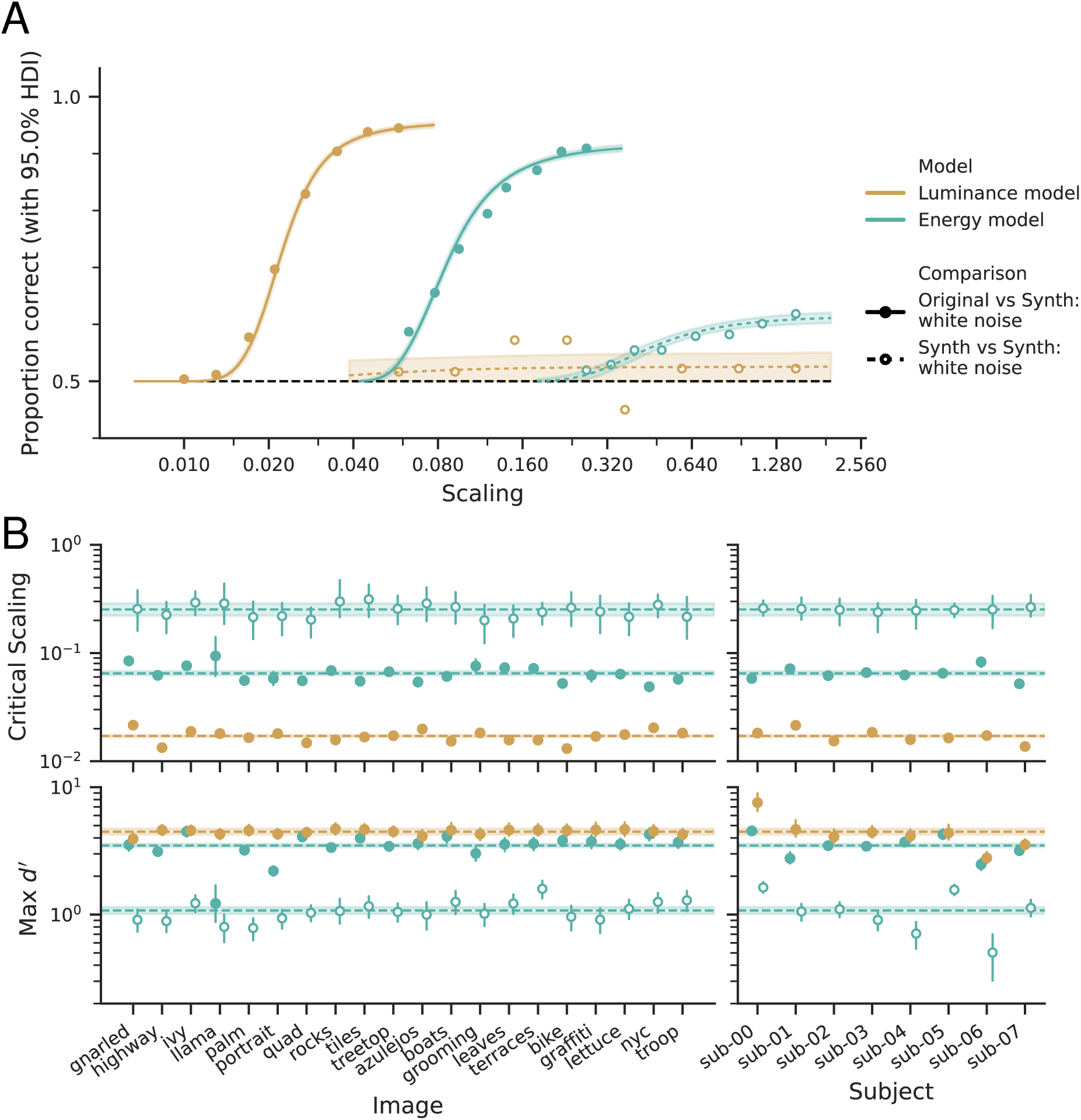
Performance curves and their corresponding parameter values for different models and stimulus comparisons. The luminance model has a substantially smaller critical scaling than the energy model, and original vs. synthesized comparisons yield smaller critical scaling values than synthesized vs. synthesized comparisons. (A) Psychometric functions, expressing probability correct as a function of scaling parameter, for both energy and luminance models (aqua and beige, respectively), and original vs. synthesized (solid line) and synthesized vs. synthesized (dashed line) comparisons. Data points represent average values across subjects and target images, 4320 trials per data point except for the luminance model synthesized vs. synthesized comparison, which has only 180 trials per data point (one subject, five target images). Lines represent the posterior predictive means of fitted curves across subjects and target images, with the shaded region indicating the 95% highest density interval (HDI, Kruschke (2015)). (B) Estimated parameter values, separated by target image (left) or subject (right). Top row shows the critical scaling value and the bottom the value of the maximum *d*^f^ parameter. Points represent the posterior means, shaded regions the 95% HDI, and horizontal dashed lines and shaded regions the global means and 95% HDI. Note that the luminance model, synthesized vs. synthesized comparison is not shown, because the data are poorly fit (panel A, beige dashed line).

### Critical scaling is smaller for original vs. synthesized comparisons than synthesized vs. synthesized comparisons

For both luminance and energy models, it is generally easier to distinguish an original image stimulus from a synthesized stimulus than to distinguish two synthesized stimuli initialized with different white noise seeds (with same target image and scaling value), as also reported in Wallis et al. (2019) for their pooled texture model. For the luminance model, discrimination of two synthesized stimuli is nearly impossible at all scaling values (note that we only have data for one participant, an author, for this comparison; we did not wish to subject the other participants to this nearly impossible task). For the energy model, discriminating two synthesized stimuli is possible but difficult, with performance only approaching 60%, on average (although note that there are substantial differences across subjects, see figure 6B and appendix 6). The critical scaling value for this comparison, approximately 0.25, is comparable to that reported in Freeman and Simoncelli (2011) for their pooled energy model. The asymptotic performance, however, is much lower in our data. We attribute this to experimental differences (see appendix 4).

The difficulty of differentiating between two synthesized stimuli is striking, as illustrated in figure 7. In the limit of global pooling windows, luminance metamers are samples of white noise, which cannot be distinguished when presented at the resolution and extent used for the stimuli in this study (Wallis et al. (2019) made a similar point when discussing their use of the original vs. synthesized task). Analogously, synthesis with the energy model forces local oriented spectral energy to match, without explicitly constraining the phase. Two instances of phase scrambling within peripheral windows are not easily discriminable, even though either of the two might be discriminable from a stimulus with more structure. The difficulty discriminating between stimulus pairs like those shown in figure 7 may arise early or late in processing, up to and including memory. Wherever this difficulty arises, it is more pronounced for the synth vs. synth comparisons. These results complicate the interpretation of the critical scaling value, which we return to in the discussion.

**Figure 7.**
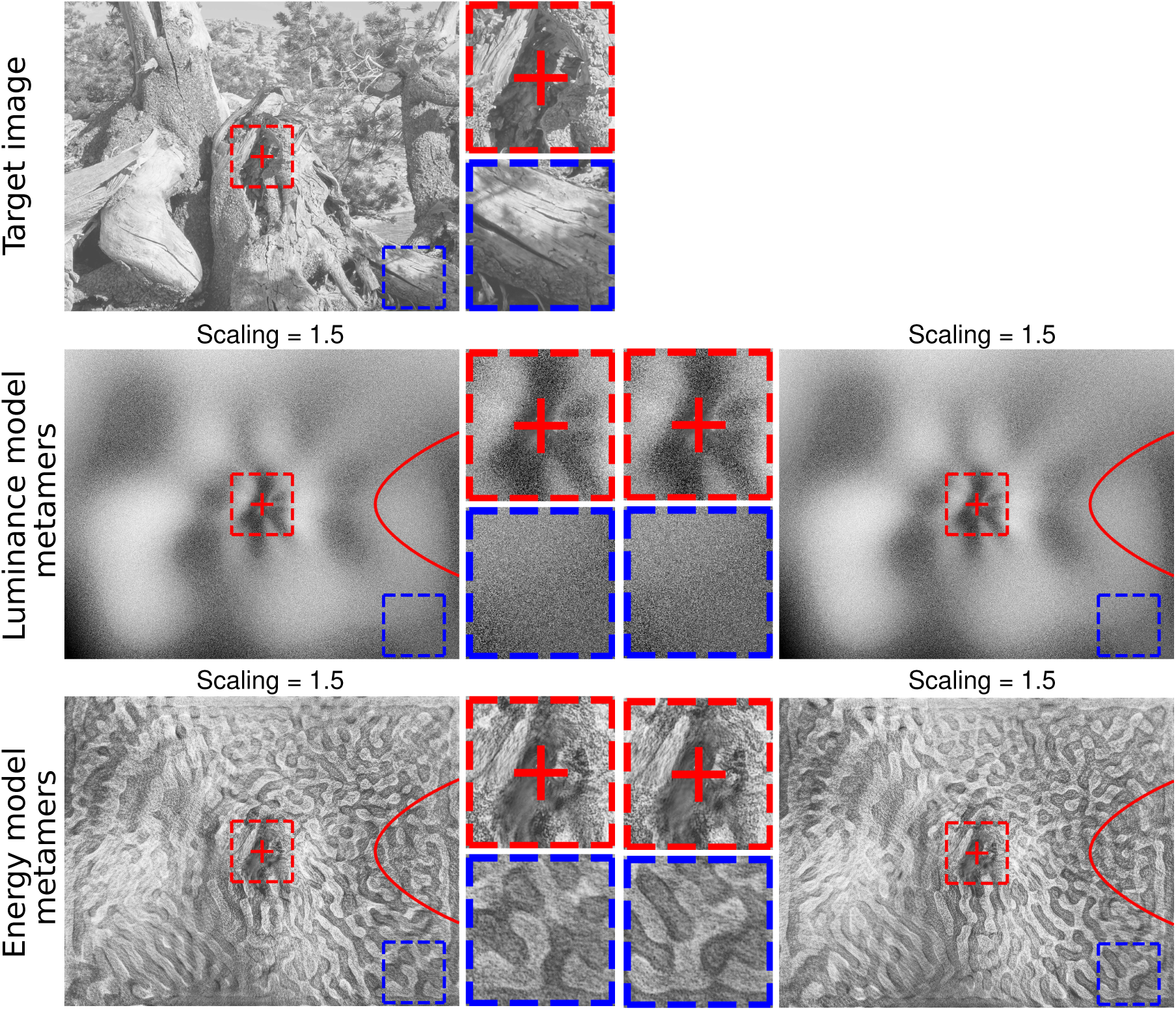
Comparison of two synthesized metamers is more difficult than comparison of a synthesized metamer with the original image. For the highest tested scaling value (1.5) the original vs. synthesized comparison is trivial while the synthesized vs. synthesized comparison is difficult (energy model) or impossible (luminance model). Top: target image. Middle: Two luminance model metamers, generated from different initial uniform noise images. Bottom: Two energy model metamers, generated from different initial uniform noise images. All four of the model metamers can be easily distinguished from the natural image at top (original vs. synthesized), but are difficult to distinguish from each other, despite the fact that their pooling windows have grown very large (synthesized vs. synthesized). Full resolution version of this figure can be found on the OSF.

### Asymptotic discrimination performance depends on target image content, but critical scaling does not

To the extent that the models (at critical scaling) capture something important about human perception, stimulus pairs that are model metamers will be perceptual metamers, and hence discrimination should be at chance. Neither model offers predictions of perceptual discriminability (they are deterministic, and do not specify any method of decoding or comparing stimuli). Consistent with this, the critical scaling, which measures the point at which stimulus pairs become indistinguishable, does not vary much across target images for a given model and comparison, unlike performance at super-threshold scaling values and the asymptotic levels of *d*^’^ (figure 6B). Variations in max *d*^’^ are especially clear in the target image-specific psychometric functions for the original vs. synthesized energy model comparison (figure 8). Specifically, for the llama target image, performance only rises slightly above chance, even at very large scaling windows. The respective target images in panel B suggest an explanation: much of the llama image is cloud-like, while the nyc image is full of sharp edges in the cardinal directions, which arise from precise alignment of phases across positions and scales. As discussed above, synthetic energy model metamers have matching local oriented spectral energy, with randomized phase information; in order to generate sharp, elongated contours for the buildings of nyc, the windows must be very small. Conversely, the appearance of the llama is captured even when the pooling windows are large. Thus, when scaling is larger than critical scaling, some comparisons become easy and some do not. However, this pattern depends on the interaction between the model’s sensitivities and the target image content; this pattern does not hold for the luminance model, or for synthesized vs. synthesized comparisons, for which both the llama and nyc target images exhibit typical performance. See appendix 6 for more details.

**Figure 8.**
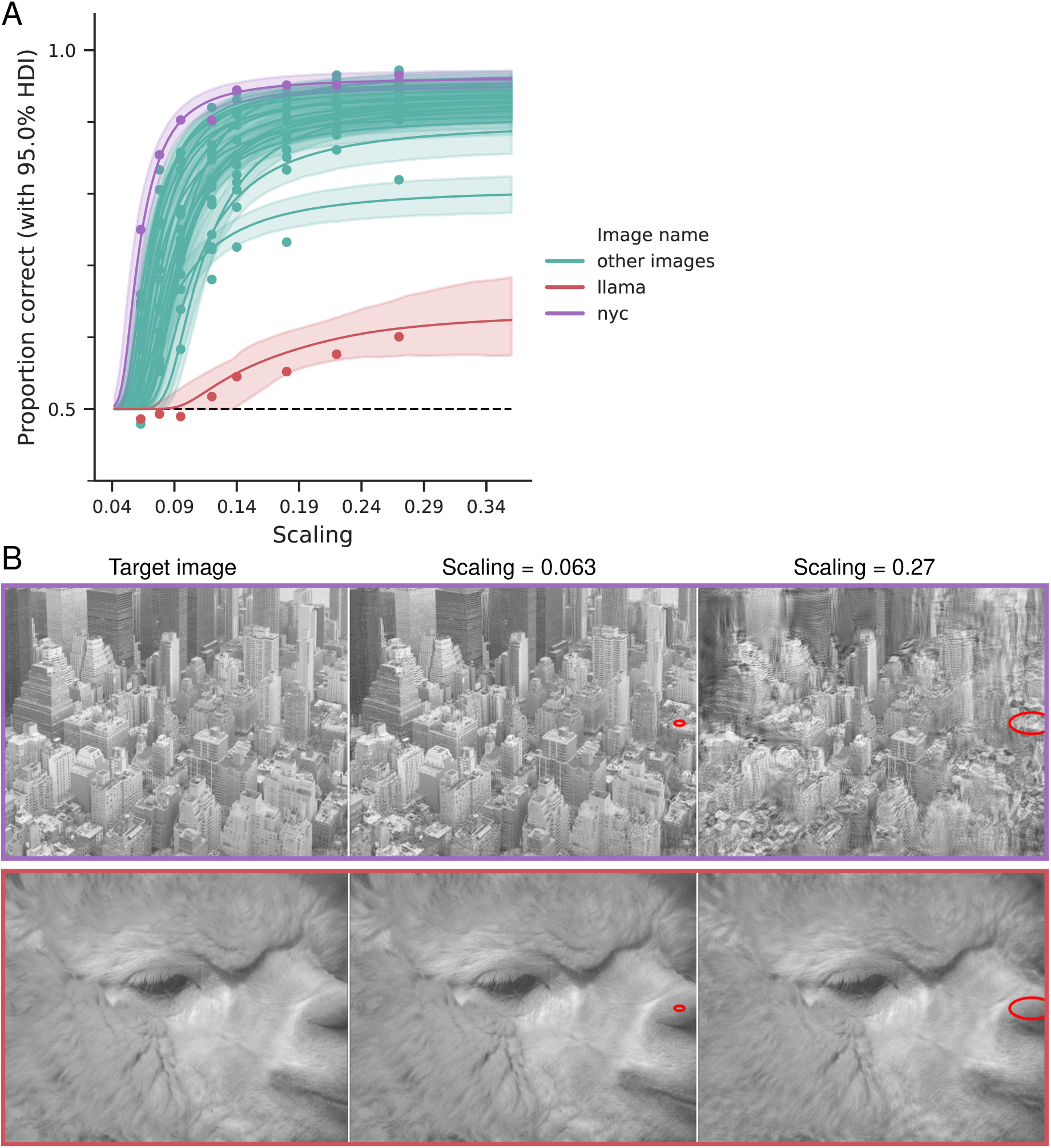
The interaction between image content and model sensitivities greatly affects asymptotic performance (especially for the synthesized vs. synthesized comparison using the energy model) while critical scaling does not vary as much. (A) Performance for each target image, averaged across subjects, comparing synthesized stimuli to natural image stimuli. Most target images show similar performance, with one obvious outlier whose performance never rises above 60%. Data points represent the average across subjects, 288 trials per data point for half the images, 144 per data point for the other half. Lines represent the posterior predictive means across subjects, with the shaded region giving the 95% HDI. (B) Example energy model metamers for two extreme target images. The top row (nyc) is the target image with the best performance (purple line in panel A), while the bottom row (llama) has the worst performance (red line in panel A). In each row, the leftmost image is the target image, and the next two show model metamers with the lowest and highest tested scaling values for this comparison. Full resolution version of this figure can be found on the OSF

We thus find evidence that although super-threshold performance depends on the interaction between target image content and model sensitivities, critical scaling is consistent across target images, in contrast with Wallis et al. (2019).

### Performance in synthesized vs. synthesized comparisons is affected by the distribution of synthesis initialization images

The two types of comparisons shown in figure 6 — original vs. synthesized and synthesized vs. synthesized — show very different critical scaling values. Specifically, for a particular scaling value and set of image statistics, some stimulus pairs are much easier to discriminate than others. We hypothesize that metamers synthesized from white noise seeds are restricted to a relatively small region of the full set of model metamers. As a result, these stimuli are more perceptually similar to each other than they are to the target stimulus. To generate metamers outside of this set, we also used other natural images from our data set to initialize the synthesis procedure (which was not done in previous studies, Freeman and Simoncelli (2011); Wallis et al. (2019)).

Figure 9A shows behavior for these additional comparisons in a single subject, sub-00. Changing the initialization image has a large effect on the synthesized vs. synthesized comparison but little-to-no effect on the original vs. synthesized comparison. For synthesized vs. synthesized, initializing with a different natural image improves performance compared to initializing with white noise, but is still worse than performance for original vs. synthesized.

**Figure 9.**
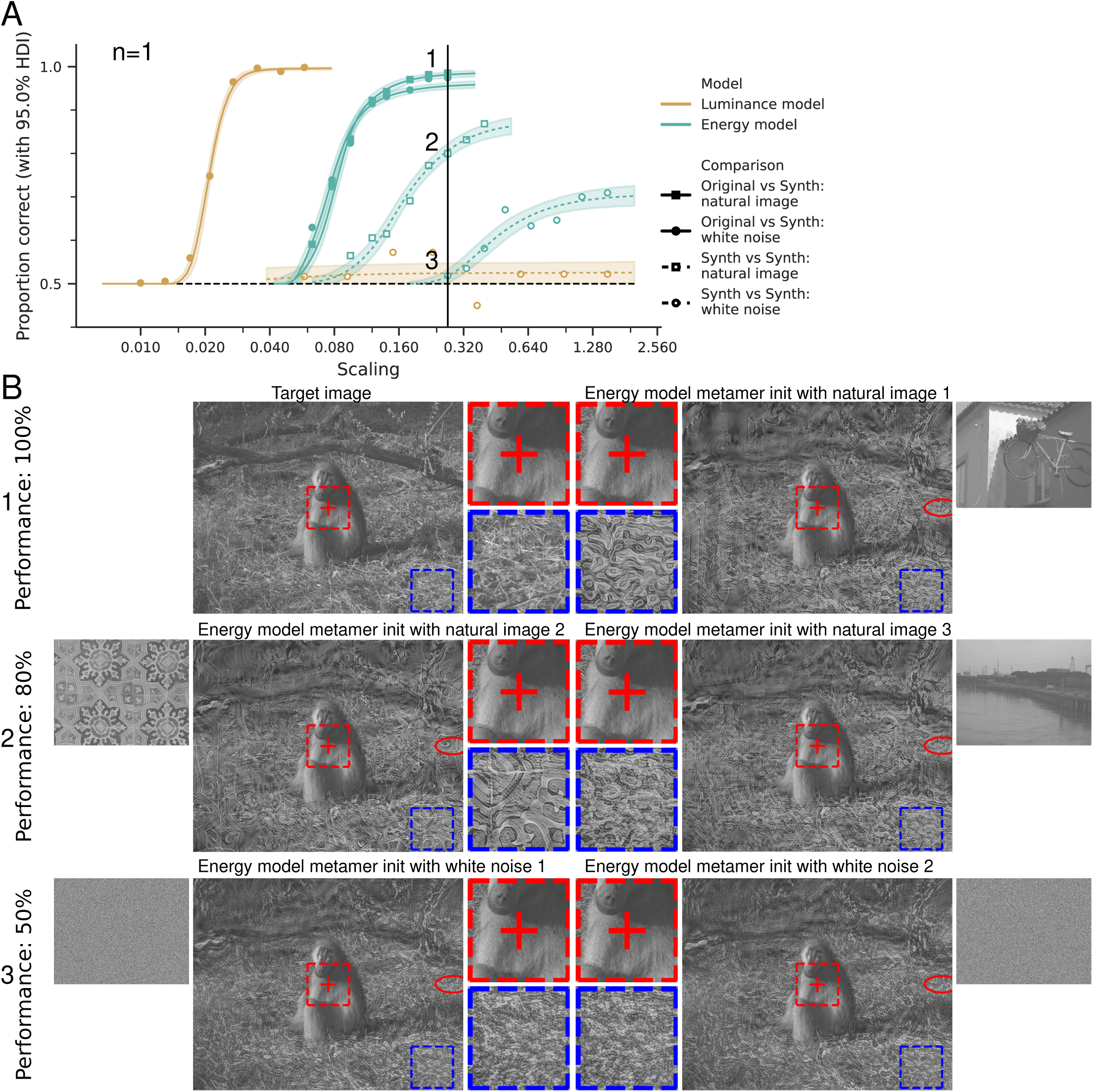
Initializing model metamers with natural images does not affect performance in the original vs. synthesized comparison, but reduces critical scaling and increases max *d*^’^ for the synthesized vs. synthesized comparison. Note all data in this figure is for a single subject and 15 of the 20 target images. (A) Probability correct for one subject (sub-00), as a function of scaling. Each point represents the average of 540 trials (over all fifteen target images), except for the synthesized vs. synthesized luminance model white noise comparison (averaged over 5 target images). Vertical black line indicates scaling value where performance ran from chance to 100%, based on initialization and comparison, as discussed in panel B. (B) Three comparisons corresponding to the three psychophysical curves intersected by the vertical black line in panel A. The small image next to each model metamer shows the image used to initialize synthesis. See text for details. Full resolution version of this figure can be found on the OSF.

**Figure 10.**
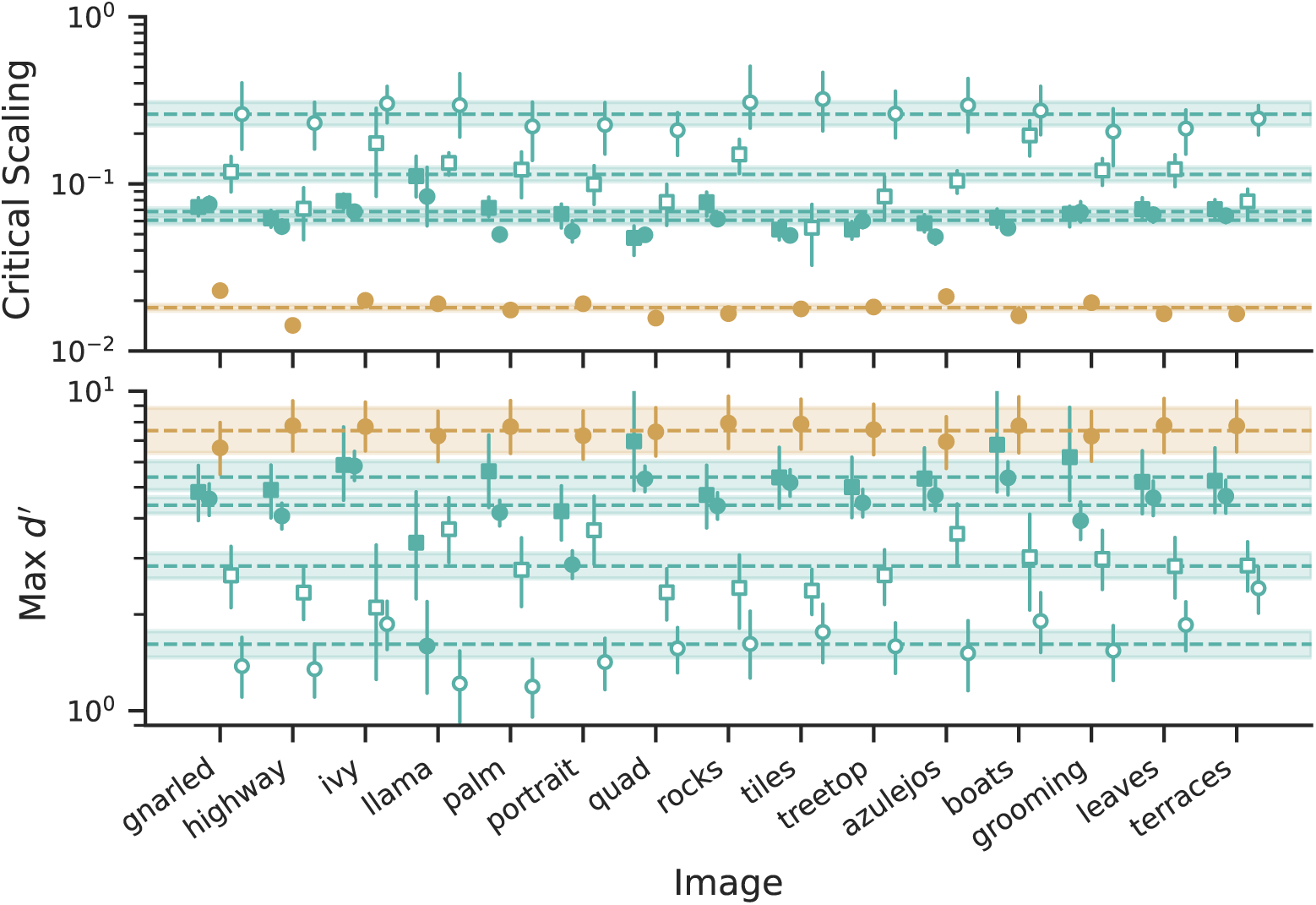
Parameter values for the comparisons shown in figure 9A (Top: critical scaling value; Bottom: max *d’*). Data shown is from the single subject who completed all comparisons. Points represent the posterior means, shaded regions the 95% HDI, and horizontal dashed lines and shaded regions average across all shown stimuli for this subject. Note that the luminance model, synthesized vs. synthesized: white noise comparison is not shown in this panel, because the data was poorly fit by this curve.

Figure 9B shows three comparisons involving five metamers arising from different initializations, each with scaling corresponding to the vertical line in panel A, but with dramatically different human performance. The top row shows the easiest comparison, between the original image stimulus and a synthesized stimulus initialized with a different natural image (bike); the subject was able to distinguish these two stimuli with near-perfect accuracy (in this case, when comparing against a natural image stimulus, performance is identical regardless of whether the metamer was initialized with white noise or natural image). The bottom row shows the hardest comparison, between two synthesized stimuli initialized with different samples of white noise. As discussed above, comparing two stimuli of this type is difficult even with large pooling windows; at this scaling level, humans are insensitive to the differences between them, and so performance was at chance. The middle row shows two synthesized stimuli, initialized with different natural images, which the subject was able to distinguish with moderate accuracy. When comparing these two stimuli, one can see features in the periphery that remain from the initial image (e.g., traces of the lines between tiles are present in the bottom right corner of the tile-initialized stimulus, while remnants of the power lines are present in the top right of the highway-initialized stimulus). Even when fixating, the subject was able to use these features to distinguish the two stimuli, i.e., the human was sensitive to them while the model was not. This reinforces the notion that the initialization of the synthesis process is important. In both the middle and bottom row, both stimuli are synthesized (i.e., neither row contains the target image) yet one comparison is much harder than the other.

Note that the experiments discussed in this section were only carried out by a single participant. We do not believe this substantially affects our conclusions. This subject was an author (and thus an expert observer, in that they were very familiar with the stimuli), and has an average critical scaling and the highest max *d*^’^ value for the comparisons we have multiple subjects’ data (see figure 6); as we largely interpret the critical scaling, values of which are quite consistent across observers, their results are therefore likely typical. While our ability to precisely estimate this critical scaling is limited by the number of subjects, we believe that the broad trends that we focus on are likely robust. That is, while the exact critical scaling values will likely vary across participants, we believe that starting from a natural image will have a negligible effect on the critical scaling of the energy model original vs. synth task, while doing so for the energy model synth vs. synth task will lead to a result intermediate between the two energy model white-noise initialization comparisons. Finally, we gathered three hours of data for these two energy model natural image comparisons, allowing us to fit the full psychophysical curve. See the methods section Session and block organization for a full enumeration of the trials.

As with the comparison (but unlike target image content), the initialization of the synthesis procedure has a substantial effect on our experimental estimates of critical scaling.

### Critical scaling is not determined by model dimensionality

For each model, the number of statistics is proportional to the number of pooling regions, and thus decreases quadratically with scaling. Table 1 shows average critical scaling values across all conditions, along with the corresponding number of model statistics. We can see that critical scaling does not correspond to a fixed number of statistics. We should also note that if one were to use the model outputs as a compressed representation of the image, the number of statistics in each representation is almost certainly an overcount, for several reasons. First, in order to ensure that the Gaussian pooling windows uniformly tile the image, the most peripheral windows in the model have the majority of their mass off the image, which is necessary to avoid synthesis artifacts. Second, for the energy model, we did not attempt to determine how the precise number of scales or orientations affected metamer synthesis, and currently all scales are equally-weighted across the image. As the human visual system is insensitive to high frequencies in the periphery and low frequencies in the fovea, this is probably unnecessary, and so some of these statistics can likely be discarded. Finally, our pooling windows are highly overlapping and thus the pooled statistics are far from independent; this redundancy means that the effective dimensionality of our model representations is less than the quoted number of statistics.

**Table 1.** Critical scaling (posterior mean over all subjects and target images) and number of statistics (as a percentage of number of input image pixels - a type of “compression rate”), for each model and comparison. Note that the critical scaling for original vs. synthesized comparisons does not result in the same number of statistics across models and, in particular, at their critical scaling values, all models have dimensionality less than that of the input image.

| Model | Comparison | Critical Scaling | Number of Statistics<br>(percentage of image pixels) |
| --- | --- | --- | --- |
| Luminance | Original vs. Synth: white noise | 0.017 | 19.6 % |
|  | Synth vs. Synth: white noise | N/A | N/A |
| Energy | Original vs. Synth: white noise | 0.065 | 34.7 % |
|  | Original vs. Synth: natural image | 0.068 | 31.5 % |
|  | Synth vs. Synth: natural image | 0.114 | 11.6 % |
|  | Synth vs. Synth: white noise | 0.252 | 2.6% |

In summary, critical scaling does not reflect a fixed number of statistics. We instead believe it reflects the spatial scale at which human perception is sensitive to changes in each of these statistics.

## Discussion

We measured perceptual discriminability of wide field-of-view metamers arising from foveated pooling of two visual attributes. We found that performance depended on the model type (i.e., the type of image features that are pooled), the nature of the comparison (original vs. synthesized, or synthesized vs. synthesized), the seed image used for metamer synthesis, and to a lesser extent, the natural image target. Specifically, critical scaling was much smaller for the luminance than for the energy model, and much smaller for original vs. synthesized than for synthesized vs. synthesized comparisons. For original vs. synthesized comparisons, critical scaling values were also reduced when the synthesis procedure was initialized from another natural image rather than white noise. Finally, asymptotic performance, but not critical scaling, was affected by target image content. Below, we consider why each of these factors affects performance.

### Why does critical scaling depend on the feature that is pooled?

Visual representations are formed through a cascade of transformations. An appealing hypothesis is that each of these is comprised of the same canonical “blurred feature extraction” computation, differing only in the choice of feature and the spatial extent of the blurring (*Fukushima, 1980*; *Douglas et al., 1989*; *LeCun et al., 1989*; *Heeger et al., 1996*; *Riesenhuber and Poggio, 1999*; *Bruna and Mallat, 2013*). In the perception literature, *Lettvin* (*1976*) provides an early, informal description of this “compulsory feature averaging” as an explanation for the degradation of peripheral percepts, non-foveated versions have been described in *Parkes et al.* (*2001*); *Pelli et al.* (*2004*); *Greenwood et al.* (*2009*), and foveated proposals appear in *Balas et al.* (*2009*); *Freeman and Simoncelli* (*2011*). In these, stages are distinguished by their features and the scaling of the pooling regions with eccentricity. Similarly, in the primate visual system, features represented in successive stages of processing become more complex and receptive fields increase in size. In particular, the optics and photoreceptors pool the incident light over small regions, V1 pools spectral energy over larger regions, V2 pools texture-related statistics over yet larger regions, and so forth. Consistent with this view, we find that for synthetic metamer stimuli, the features pooled by the model have a dramatic effect on the critical scaling value, which is approximately four times larger for the energy model than for the luminance model in the original vs. synthesized comparison (figure 6 and table 1). The values of critical scaling as a function of image statistic and comparison type are summarized in figure 11, along with the results of *Freeman and Simoncelli* (*2011*) and *Wallis et al. (2019)* obtained for a model of texture (*Portilla and Simoncelli, 2000*). Overall, critical scaling for original vs. synthetic comparisons follows approximate ratios of 1:4:12 for luminance:spectral-energy:texture features (filled circles, figure 11).

**Figure 11.**
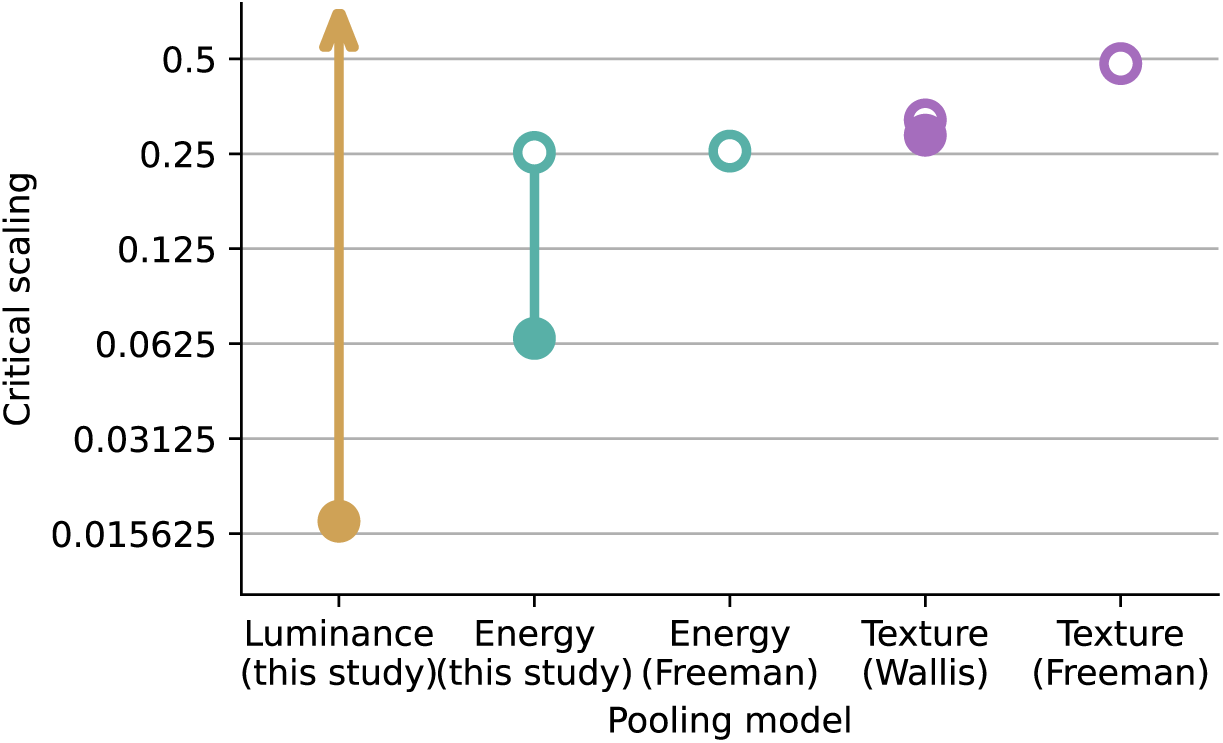
Critical scaling values for the two pooling models presented in this paper (Luminance and Energy) along with a Texture model (data from Freeman and Simoncelli (2011) and Wallis et al. (2019), averaging across the two image classes). Filled points indicate original vs. synthesized comparisons, while unfilled points indicate synthesized vs. synthesized comparisons (for the luminance model, this is effectively infinite, since participants were unable to perform the task for any scaling value). For all three models, critical values are smaller in the original vs. synthesized comparison than the synthesized vs. synthesized one, but this effect decreases with increasing complexity of image statistics. Our critical scaling values for synth vs. synth comparisons of the energy model are consistent with those reported by Freeman and Simoncelli (2011).

### Why does critical scaling depend on comparison type?

We found large effects of comparison type on performance (figure 11, filled vs. unfilled). Specifically, for the energy model the critical scaling for synthesized vs. synthesized was about four times larger than that for original vs. synthesized. For the luminance model, participants were generally unable to discriminate any pairs for the synthesized vs. synthesized comparison. These effects arise from an interaction between human perception, the model, and the synthesis process.

As shown in figure 12, the synthesis procedure can bias the sampling of model metamers such that two synthesized stimuli are metameric with each other, but distinguishable from the target stimulus. The idealized version of the metamer paradigm implicitly assumes that the synthesized stimuli sample the manifold of possible model metamers broadly, but our synthesis procedure (as with that of Freeman and Simoncelli (2011) and Wallis et al. (2019)) does not guarantee this.

**Figure 12.**
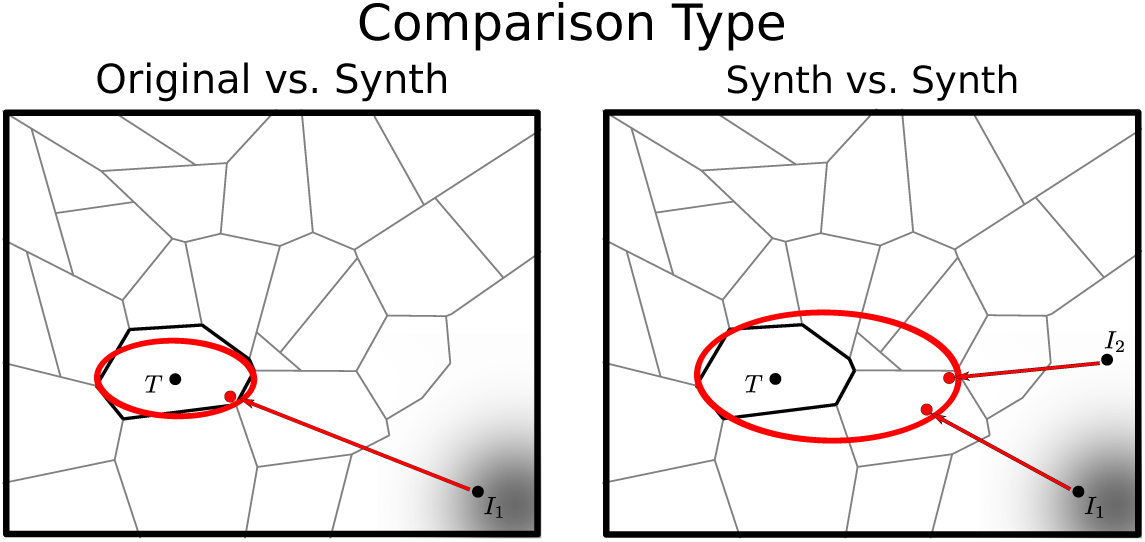
Illustration of how critical scaling value can depend on comparison type. Left: For the original vs. synthetic comparison, critical scaling corresponds to the largest ellipse such that the synthetic image is indistinguishable from (i.e., lies within the same perceptual region as) the target (original) image *T*. Right: Example configuration in which synthetic vs. synthetic comparisons lead to a larger critical scaling (with correspondingly larger ellipse). The two synthesized images are indistinguishable from each other, but not from the target.

The conceptual framework we outlined in this paper emphasizes deterministic information loss, arising from projections of high-dimensional visual images onto a lower-dimensional manifold. Under these conditions, metamerism is transitive: all stimuli within an equivalence class are indistinguishable from each other, and all stimulus pairs from different classes are potentially distinguishable. However, reductions in discriminability can also arise from stochastic information loss: discrimination thresholds are governed by noisy neural responses, as developed in the frameworks of signal detection theory and ideal observer models. The psychophysical function we fit to the data arises from an observer model that compares local statistics of stimuli (Freeman and Simoncelli, 2011), and combines the two forms of information loss. It asserts that performance falls *exactly* to chance when the scaling used for synthesis is less than or equal to the true scaling (i.e., when the stimuli are metameric, projecting to the identical point in perceptual space). As the synthesis scaling exceeds the true scaling, the measured differences in stimulus statistics exceed the level of noise in subjects’ perceptual systems, and performance gradually improves. Our analysis thus focuses on the critical scaling, the point at which the piecewise function first rises above chance, which is the primary prediction of the metamer paradigm. In practice, our metamer models, like all models (Box, 1976), are imperfect, but small errors in the models are likely masked by subjects’ stochastic noise during discrimination.

### Why does this comparison-type dependency decrease with feature complexity?

The effect of comparison type was previously reported by Wallis et al. (2019) and Deza et al. (2019) in models pooling texture-like statistics. As seen in figure 11, the ratio in critical scaling between the two types of comparisons decreases as the model features become more complex: infinite for luminance, roughly quadruple for energy, and less than double for texture. One potential explanation for this observation is that these computations are being performed in deeper stages of the visual hierarchy and there are progressively fewer opportunities to discard information later in the hierarchy. For example, the difference in V1 responses for a pair of stimuli may be discarded in later stages, whereas in a later area (e.g., area IT), there are fewer remaining stages of processing in which information can be discarded. This may explain why we see no overlap between the critical scaling values for original vs. synthesized and synthesized vs. synthesized comparisons across target images in figure 6B, whereas Wallis et al. (2019) find substantial overlap for the texture model.

Another potential explanation is schematized in figure 13: for the luminance model, the model metamer classes grow with scaling *s* in such a way that model metamers generated from white noise initialization fall within the same perceptual metamer class for a wide range of *s* values. This is an extreme case of the synthesis issue described in the previous section. Conversely, as the texture model’s critical scaling dependence on comparison type is much weaker, we can hypothesize that its metamer classes are such that model metamers synthesized from white noise readily fall into distinct perceptual metamer classes, leading to a critical scaling value that is similar to the value for the original vs. synth comparison.

**Figure 13.**
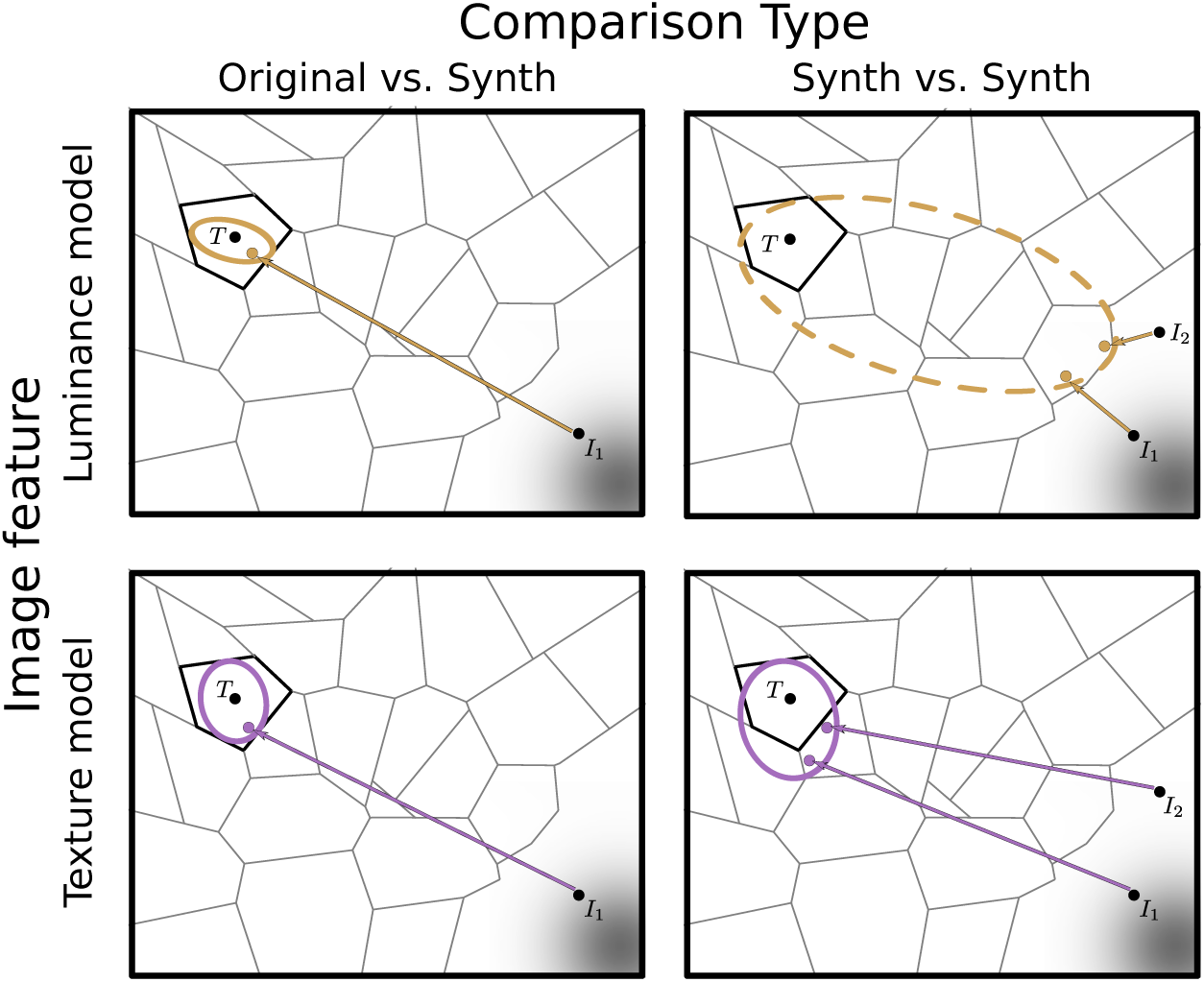
Illustration of how discrepancies between the original vs. synth and synth vs. synth comparisons can be model-dependent. Upper panels depict a hypothetical luminance model, for which pairs of synthesized stimuli are indistinguishable for all scaling values (i.e., they lie within the same perceptual metamer class for both the small ellipse on the left, and for larger ellipse on the right). In this case, critical scaling cannot be estimated (indicated by dashes in ellipse). Lower panels depict a hypothetical texture model, for which synth vs. synth comparisons yield a critical scaling value (right) only slightly larger than that obtained for original vs. synth (left). Energy model metamers (not shown) lie between these two extremes (see figure 11).

### Why does critical scaling sometimes depend on synthesis initialization?

Our experimental results demonstrate that critical scaling depends on the comparison type and feature complexity. We also find that it depends on synthesis initialization. Specifically, the set of metamer stimuli generated for our experiments is a biased sample from the space of all possible model metamers, and is particularly dependent on the distribution of initialization images. This effect is especially noticeable in the case of the luminance model, as discussed in the previous section. We conjectured that natural images are more likely than synthetic images to include information that the human visual system has evolved to discriminate and hence doesn’t discard even in later processing. As such, initialization with natural images can provide an intuitive method of sampling from a different portion of the manifold of possible model metamers. The schematic presented in figure 14 provides an illustration of how this might occur.

**Figure 14.**
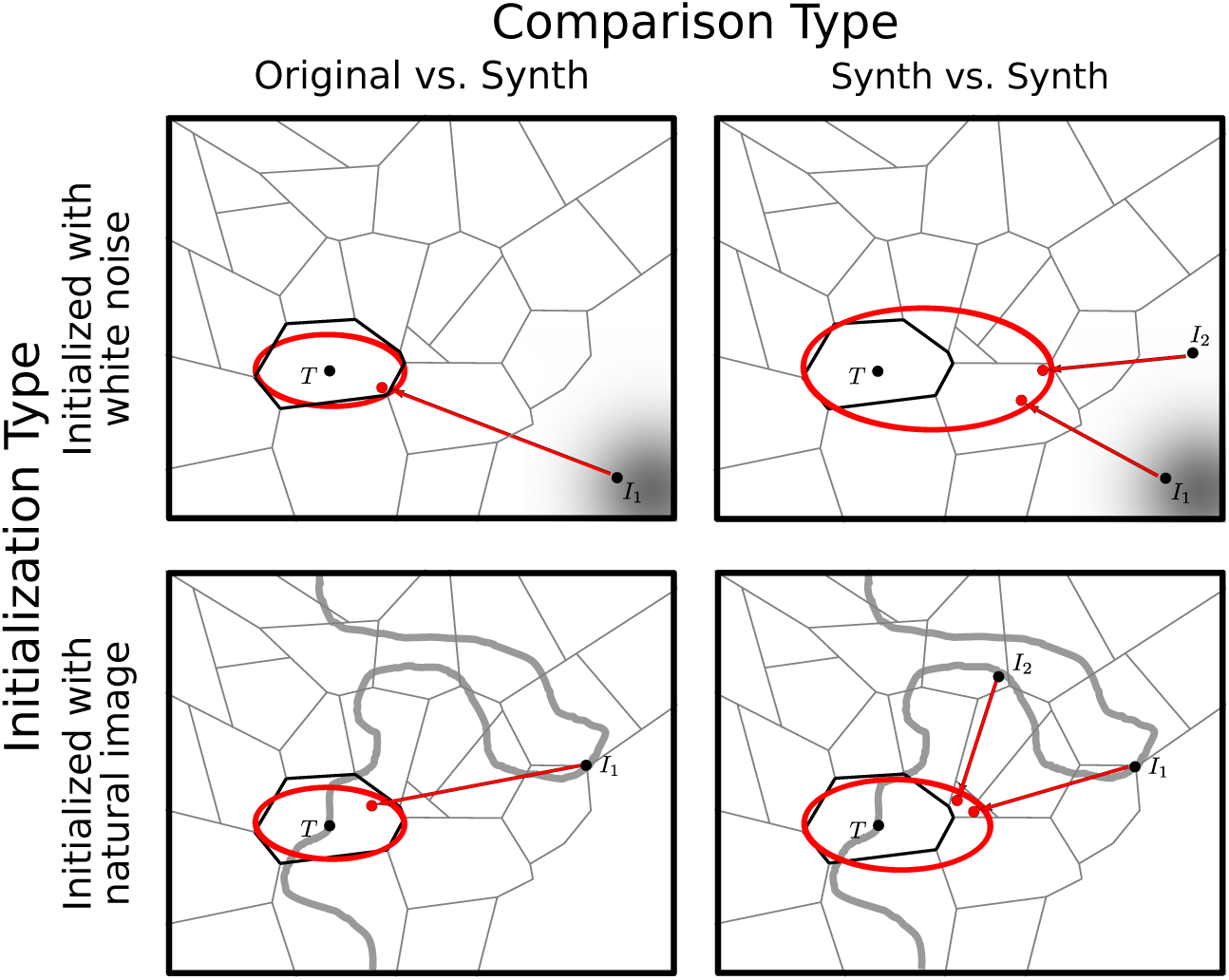
Illustration of how discrepancies between original vs. synth and synth vs. synth comparisons can depend on the image used to initialize synthesis. Initializing with white noise (as is commonly done) can lead to a large difference between the two comparisons (see also figures 12 and 13). Initializing with a natural image can reduce the magnitude of the difference.

**Figure 15.**
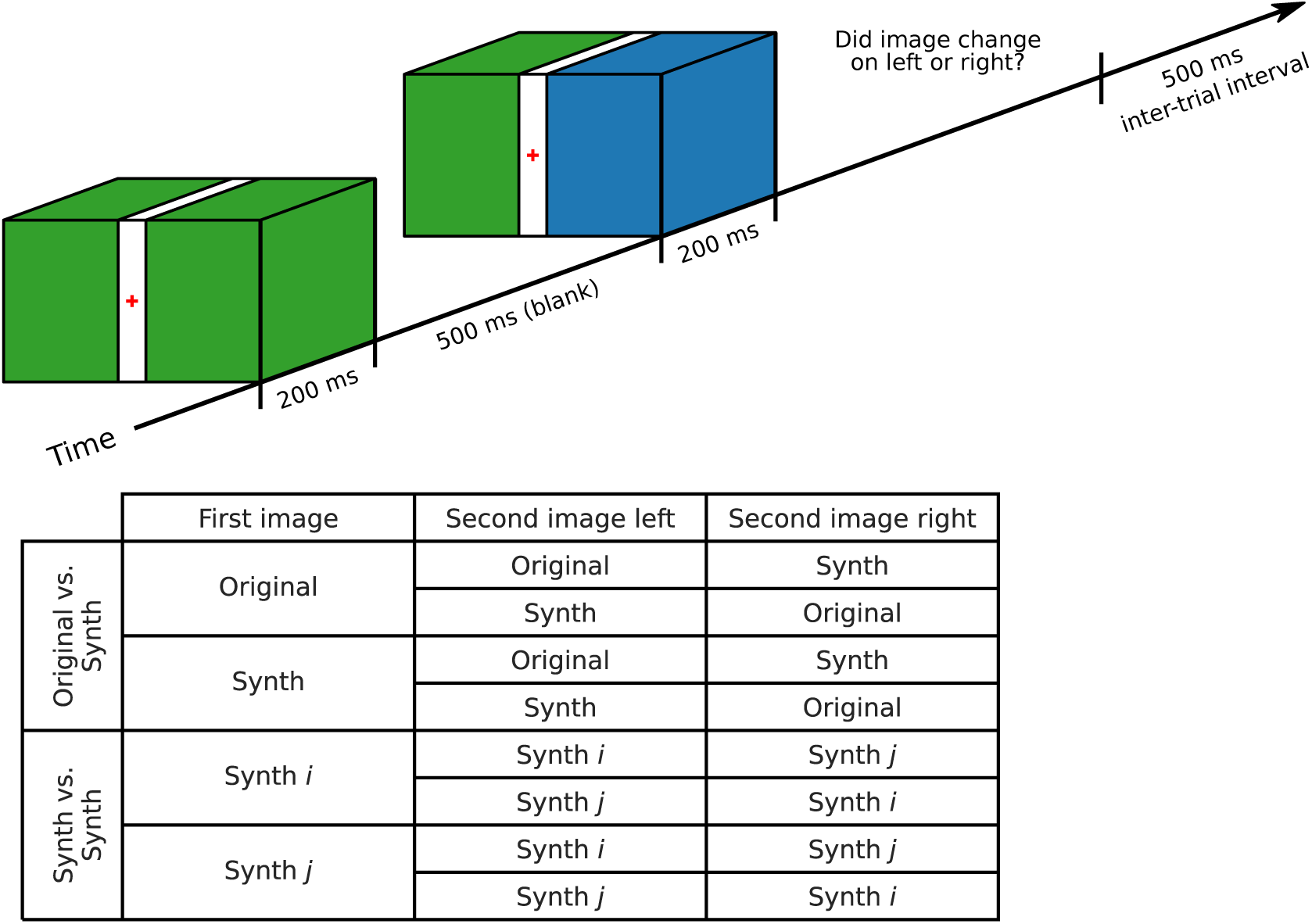
Schematic of psychophysics task. Top shows the structure for a single trial: a single stimulus is presented for 200 msec, bisected in the center by a gray bar, followed by a blank screen for 500 msec. The stimulus is re-displayed, with a random half of the stimulus changed to the comparison stimulus, for 200 msec. The participants then have as long as needed to say which half of the stimulus changed, followed by a 500 msec intertrial interval. Bottom table shows possible comparisons. In original vs. synthesized, one stimulus was the target image stimulus whose model representation the synthesized stimuli match (see figure 5), and the other was one such synthesized stimulus. In synthesized vs. synthesized, both were synthesized stimuli targeting the same target image stimulus, with the same model scaling, but different initialization. In experiments, dividing bar, blanks, and backgrounds were all midgray. For more details see text.

We controlled the generation of metamer stimuli in our experiments by selecting one of two initialization distributions: white noise and natural images. A natural intermediate choice would be pink noise, with frequency spectrum matched to that of natural images. Such initialization would reduce the high frequencies found in the super-threshold luminance model metamers (e.g., figure 4), while avoiding the detectable features found in those initialized with natural images (see appendix 2 for illustrative example images). This would result in another psychophysical curve on figure 9A, shifted to the right of the leftmost (Original vs. Synth: white noise) curve for this model, and thus a higher estimated critical scaling value. The logic of our experiments, however, relies on finding the *smallest* critical scaling value across initializations, rather than generating stimuli that look as similar to the target as possible. This reiterates the importance of thinking carefully about synthesis initialization for metameric stimulus generation, but we expect such initializations would not otherwise influence our interpretation of the results. A more principled statistical sampling approach could result in metamers that better reflect natural image properties, or metamers that are more likely to be discriminable by human observers.

### Why does asymptotic performance, but not critical scaling, depend on image content?

Similar to Wallis et al. (2019) and Brown et al. (2023), we find that metamer discrimination performance is somewhat dependent on image content. Both of those studies synthesize model metamers based on pooled texture statistics, and Wallis et al. (2019) shows that texture-like target image stimuli are harder to distinguish from their synthesized stimuli than scene-like ones, while Brown et al. (2023) shows that original textures with higher global and local regularity (e.g., woven baskets) are easier to distinguish from their synthesized stimuli than those with low regularity (e.g., animal fur). This aligns with our result: the most distinguishable pairs include natural image stimuli features not well-captured by the synthesizing model, whereas the least distinguishable include those natural image stimuli whose features are all adequately captured.

However, we should note that we found this image-level variability largely in super-threshold performance, and this variability does not constitute a failure of these pooling models. As pointed out by Freeman and Simoncelli (2011), asymptotic performance also varies with experimental manipulation, while critical scaling remains relatively unaffected. The metamer paradigm makes strong predictions about what happens when the representations of two stimuli are matched: they are indistinguishable, and so performance on a discrimination task will be at chance, as captured by the critical scaling value. However, it makes *no* predictions about performance at super-threshold levels. An analogy with color vision seems apt: color matching experiments provide evidence for what spectral distributions of light are perceived as identical colors, but provide no information about whether humans consider two shades of blue distinguishable or blue more similar to green or to red; further measurements are necessary to understand color appearance.

By investigating how such differences are handled by later brain areas, this complementary approach could gain a better understanding of stimulus discriminability beyond “identical or not”. Such work could use the models presented here as a starting point and could draw on the substantial literature of observer models in vision science and image processing; see Ziemba and Simoncelli (2021); Kurzawski et al. (2024) for examples applying this approach to pooling model metamer images. Combining and extending the synthesis-focused metamer approach with observer models’ attention to super-threshold performance would help lead to a fuller understanding of human perceptual sensitivities and insensitivities.

## Conclusion

In summary, we’ve used large field-of-view stimuli matched to the foveation properties of the human visual system in order to estimate the spatial precision of the perceptual representation of low-level image features. By improving our understanding of the information that is discarded, these measurements provide a starting point for building a foveated model of the visual system’s front end. We’ve also provided a unified explanation of the variations in critical scaling that arise from changes in image content, comparison type, and synthesis initialization, providing guidance on how to use model and perceptual metamers for understanding visual representations.

## Materials and Methods

All experimental materials, data, and code for this project are available online under the MIT or similarly permissive licenses. Specifically, software is on GitHub, synthesized metamers can be browsed on this website, and all stimuli and data can be downloaded from the OSF. The GitHub site provides instructions for downloading and using data.

### Synthesis

We synthesized model metamers matching 20 different natural images (the target images) from the authors’ personal collections, as well as from the UPenn Natural Image Database (Tkačik et al. (2011), extended dataset of urban images provided by David Brainard). The selected photos were high-resolution with 16-bit pixel intensities proportional to luminance, that had not undergone lossy compression (which could result in artifacts). They were converted to grayscale using scikit-image’s color.rgb2gray function (van der Walt et al., 2014), cropped to 2048 by 2600 pixels (the Brainard photos were 2014 pixels tall, so a small amount of reflection padding was used to reach 2048 pixels), and had their pixel values rescaled to lie between 0.05 and 0.95. Synthesized stimuli were still allowed to have pixel values between 0 and 1; without rescaling the target images, synthesis resulted in strange artifacts with pixels near 0, as this was the minimum allowed value. The target images were chosen to span a variety of natural image content types, including buildings, animals, and natural textures (see figure 5).

We synthesized the model metamers using custom software written in PyTorch (Paszke et al., 2019), using the AMSGrad variant of the Adam optimization algorithm (Kingma and Ba, 2014; Reddi et al., 2018), with learning rate 0.01. Slightly different approaches were used for the luminance and energy model metamers. For the luminance model metamers, the objective function was to minimize the mean-squared error between the model representation of the target and synthesized stimuli, *L(x,x^) = (M(x) − M(x^))^2^*, and synthesis was run for 5000 iterations. For the energy model metamers, the objective function also contained a quadratic range penalty term, which penalized any pixel values outside of [0, 1], *L(x,x^) = 0.5(M(x) − M(x^))^2^ + 0.5B(x^)*, where

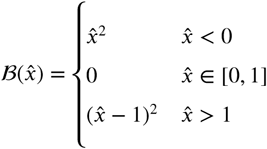

Synthesis was run for 15000 iterations. Additionally, energy model metamer synthesis used stochastic weight averaging (Izmailov et al., 2018), which helped avoid local optima by averaging over pixel values as synthesis neared convergence, and used coarse-to-fine optimization (Portilla and Simoncelli, 2000). Additionally, each statistic (in both models) was z-scored using the average statistic value computed across the entire image on a selection of grayscale texture images. For both models, synthesis terminated early if the loss had not decreased by more than 10^-9^ over the past 50 iterations. While not all model metamers achieved the same loss values, with differences in synthesis loss across target images, there was no relationship between the remaining loss and behavioral performance.

For each model, its windows were represented as two tensors, one for angular slices and one for annuli, which, when multiplied together, would give the individual windows, with separate sets of windows for each scale in the energy model. This required a large amount of memory, and so for scaling values below 0.09, models were too large to perform synthesis on the available NVIDIA GPUs with 32GB of memory. Thus, all luminance model metamers were computed on the CPU, and synthesis of a single stimulus took from about an hour for scaling 1.5 to 2 days for scaling 0.058 to 14 days for scaling 0.01. For the energy model metamers, the lowest two scaling values were computed on the CPU, with synthesis taking about a week. For those energy model metamers which were able to be computed on the GPU, synthesis took from 5 hours for scaling 0.095 to 1.5 hours for scaling 0.27 and above. This synthesis procedure was completed in parallel using the high-performance computing cluster at the Flatiron Institute.

Synthesized stimuli for original vs. synthesized and synthesized vs. synthesized white noise comparisons (see Psychophysical experiment) were initialized with full-field patches of white noise (each pixel sampled from a uniform distribution between 0 and 1). For each model, scaling value, and target image, three different initialization seeds were used. A unique set of three seeds was used for each scaling value and target image, except for the following, which all used seeds {0, 1, 2}:

- Luminance model: azulejos, bike, graffiti, llama, terraces, tiles; scaling 0.01, 0.013, 0.017, 0.021, 0.027, 0.035, 0.045, 0.058, 0.075 and 0.5.
- Energy model: azulejos, bike, graffiti, llama, terraces, tiles; scaling 0.095, 0.12, 0.14, 0.18, 0.22, 0.27, 0.33, 0.4, and 0.5.

For original vs. synthesized and synthesized vs. synthesized natural image comparison, synthesized stimuli for each model, scaling value, and target image were initialized with three random choices from among the rest of the target images.

### Pooling windows

Pooling model windows are laid out in a log-polar grid, with peaks spaced one standard deviation apart, such that adjacent window functions cross at a value of 0.352 (relative to max of 0.4). They have a single parameter, scaling, which specifies the ratio of the pooling window diameter at fullwidth half-max in the radial direction and the distance of its center from the fovea, both in degrees. For example, the pooling windows of a model with scaling factor 0.1 have a radial diameter of 1 degree at 10 degrees eccentricity, 2 at 20 degrees, etc.

Stimulus pixels within 0.5 degree from the fixation point were exactly matched in our synthesized stimuli, approximating the fovea, where no pooling occurs. Additionally, for small scaling values, windows for some distance beyond this region would be smaller than a pixel and so the only solution is to match the pixel values in that region directly. For example, with image resolution of 2048 by 2600 and display size of 53.6 by 42.2 degrees, models with scaling value of 0.063 have windows whose diameter at FWHM is smaller than a pixel out to 0.52 degrees, with this number increasing quadratically as scaling decreases, reaching 3.29 degrees for scaling 0.01 (see appendix 5 for how mean-squared error varied across eccentricity).

### Observers

Eight people (5 women and 3 men, aged 24 to 33), including an author (W.F.B.), participated in the study and were recruited from New York University. All subjects were graduate students or postdocs in vision science labs who had experience participating in other visual psychophysics experiments, and who were accustomed to performing peripheral tasks while maintaining central fixation. All had normal or corrected-to-normal vision. Each subject completed nine one-hour sessions. One subject (sub-00, author W.F.B.) also performed seven additional sessions. All subjects provided informed consent before participating in the study. The experiment was conducted in accordance with the Declaration of Helsinki and was approved by the New York University ethics committee on activities involving human subjects.

### Psychophysical experiment

A psychophysical experiment was run in order to determine which of the synthesized model metamers were also perceptual metamers. We first describe the structure of a single trial, then how the trials were organized into blocks and sessions.

### Trial structure

See figure 15 for schematic. Observers viewed a series of grayscale image stimuli on a monitor, at a size of 53.6 by 42.2 degrees. An initial stimulus, divided in half by a vertical midgray bar 2 degrees wide, was displayed for 200 msecs, before being replaced by a midgray screen for 500 msecs, followed by a second stimulus for another 200 msecs (also divided by a vertical midgray bar). Stimuli were presented for 200 msecs to minimize the possibility of eye movements. The dividing bar prevented participants’ use of discontinuities between the two stimulus halves to perform the task. One side of the second stimulus (left half or right half) was identical to the first stimulus, and the other side changed. After the second stimulus was viewed, a midgray screen appeared with text prompting a response, and the observer’s task was to report which half had changed. The observer had as much time as necessary to respond. The two compared stimuli were either two synthesized stimuli (synthesized for identical models with the same scaling value and target image, but different initializations) or one synthesized stimulus and its target image stimulus. Either stimulus could be presented first.

The midgray blank screen presented between the two stimulus presentations reduces motion cues participants could use to discriminate the two stimuli. Our models aim to capture the steady state response to the stimuli, not the transient response. The mask forces the participants to use the image content to discriminate between the two stimuli, rather than relying on temporal edges (analogous to our use of the vertical bar to prevent the use of spatial edges). This introduces a shortterm memory component in the task (participants must remember the first stimulus in order to compare it to the second stimulus), as in previous metamer discrimination experiments (Freeman and Simoncelli, 2011; Deza et al., 2019; Wallis et al., 2019). We believe the precise duration of this mask is unimportant for our results: first, Bennett and Cortese (1996) found the duration of a blank screen did not affect thresholds in a spatial frequency discrimination task over a range from 200 to 10,000 msec, and second, mask duration is likely to have a similar effect on performance as stimulus presentation duration, which Freeman and Simoncelli (2011) found affected asymptotic performance but not critical scaling.

### Session and block organization

Across 9 sessions, each subject completed a total of 12,960 trials, factored into 3 model/comparison combinations by 8 scaling values by 15 target images by 36 repetitions. This large number of trials enabled us to check individual differences within both target images and observers for each model/comparison combination. The 3 model/comparison combinations were 1) luminance model, original vs. synth, white noise; 2) energy model, original vs. synth, white noise; and 3) energy model, synth vs. synth, white noise. The 8 scaling values were logarithmically spaced, with the range chosen separately for each model/comparison to span an appropriate range of values. For luminance model, original vs. synth, white noise, the scaling endpoints were 0.01 and 0.058; for energy model, original vs. synth, white noise, the endpoints were 0.063 and 0.27; and energy model, synth vs. synth, white noise, the endpoints were 0.27 and 1.5. There were a total of 20 target images, but each subject only saw 15. Every subject saw images 1 through 10. Half the subjects also saw images 11 through 15, and half saw images 16 through 20 (see figure 5). The 36 repetitions were averaged for analysis and included 12 trials for each of 3 synthesis seeds. For the white noise-initialized comparisons, these seeds were independent samples of white noise used to initialize the synthesis procedure, resulting in 3 distinct model metamers.

Each of the above model/comparisons was tested across 3 sessions, each lasting approximately one hour. Each subject started with either the luminance or energy model, original vs. synth, white noise. The 3 sessions required for the model/comparison tested first were completed before moving onto 3 sessions testing the other model. The order of the two models was randomized across subjects. After completing these 6 sessions, the subject completed 3 sessions testing the energy model, synth vs. synth, white noise. This comparison was last as it was the most difficult.

Each of the 9 sessions consisted of 1,440 trials, containing all 36 repetitions for all 8 scaling values for 5 of the 15 target images viewed by the subject (target images were randomly assigned to sessions, independently for each subject). The 1,440 trials per session were broken up into 5 blocks of 288 trials each. Each block took about 8 to 12 minutes, and consisted of 12 repetitions for all 8 scaling values for 3 of the 5 target images.

In addition, one subject (sub-00) completed 7 additional sessions (10,080 additional trials). This included 1 session for luminance model, synth vs. synth, white noise; 3 for energy model, original vs. synth, natural image; and 3 for energy model, synth vs. synth, natural image. As with the comparisons that all subjects completed, these sessions each included 1,440 trials, factored into 5 target images by 8 scaling values by 36 repetitions. Only 1 session was included for luminance model, synth vs. synth, white noise because performance was at chance for all target images and all scaling values (see figures 6 and 7). No sessions were completed for the luminance model, natural image comparisons due to the time required for synthesis; see appendix 1 for more information.

The four types of comparisons are explained in full below:

1. Original vs. synthesized, white noise: the two stimuli being compared were always one synthesized stimulus and its target stimulus, and the synthesized stimulus was initialized with a sample of white noise.
2. Synthesized vs. synthesized, white noise: both stimuli were synthesized, with the same model, scaling value, and target stimulus, but different white noise seeds as synthesis initialization.
3. Original vs. synthesized, natural image: the two stimuli being compared were always one synthesized stimulus and its target stimulus, and the synthesized stimulus was initialized with a different natural image drawn randomly from our set.
4. Synthesized vs. synthesized, natural image: both stimuli were synthesized, with the same model, scaling value, and target stimulus, but initialized with different natural images from our set.

Subjects completed several training blocks. Before their first session, they completed an initial training block, comparing two natural image stimuli and two noise samples (one white, one pink). Before their first session of each comparison type including a natural image stimulus, they completed a secondary training block showing two natural image stimuli and two synthesized stimuli of the type included in the session, one with the largest scaling included in the task and one with the smallest. Before the session comparing two synthesized stimuli, they similarly completed a training block comparing four synthesized stimuli, two with a low scaling value and two with a high scaling value, for each of two target images. Each training block took one to two minutes and was repeated if performance on the high scaling synthesized stimuli was below 90% or subjects expressed uncertainty about their ability to perform the task (participants were expected to perform close to chance for the low scaling synthesized stimuli). Additionally, before each session which included a natural image stimulus (the original vs. synthesized comparisons), subjects were shown the five natural images that would be part of that session, as well as two example synthesized stimuli per target image, one with a low scaling value, one with a high scaling value. Before each session comparing two synthesized stimuli (the synthesized vs. synthesized comparison), subjects were shown four example synthesized stimuli per target image, two with the lowest scaling value and two with the highest scaling value for that comparison. A video of a single energy model training block, original vs. synthesized: white noise comparison, can be found on the OSF.

### Apparatus

The stimuli were displayed on an Eizo CS2740 LED flat monitor running at 60 Hz with a resolution of 3840 by 2160 pixels. The monitor was gamma-corrected to yield a linear relationship between luminance and pixel value. The maximum, minimum, and mean luminances were 147.73, 0.3939, and 77.31 cd/m^2^, respectively.

The experiment was run with a viewing distance of 40 cm, giving 48.5 pixels per degree of visual angle. A chin and forehead rest was used to maintain head position, but the subjects’ eyes were not tracked.

The experiment was run using custom code written in Python 3.7.0 using PsychoPy 3.1.5 (Peirce et al., 2019), run on an Ubuntu 20.04 LTS desktop. A button box was used to record the psychophysical response data. All stimuli were presented as 8-bit grayscale images.

A limitation of the study is that we did not track gaze direction. However, there are several reasons to think that fixation breaks do not strongly influence our conclusions. First, the two-degree wide vertical bar at the center of the screen ensures that minor deviations in gaze direction from the fixation point would have little effect. Second, each stimulus was only presented for 200 msecs, which is the typical latency of saccades (Carpenter, 1988). This does not rule out saccades during the trials, but participants did not have the time to freely explore the images. In principle, participants could have begun the trials gazing away from the fixation point. However, all of our participants were trained observers with experience doing peripheral tasks while maintaining central fixation. Moreover, variation in fixation across observers would likely reveal itself in variability in estimated psychometric curves. But the consistency of estimated parameters (indicated both by their relatively narrow credible intervals and similarity to results reported in Freeman and Simoncelli (2011)) suggests that any such variation did not greatly affect our results. A participant could cheat on the task by, for example, consistently fixating in the corner of the screen. However, such a subject would be an outlier, which we do not see in our data. Indeed, the participant who is an author, and was thus highly motivated to maintain fixation, did not have worse performance than the other observers. Finally, our interpretation of our results relies on the ordering of the critical scaling across models and comparisons, rather than their precise values. Given the consistency of these estimates and their separation from each other, we do not believe the lack of eye tracking affects the overall pattern of results, and hence our conclusions.

### Data analysis

All trials were analyzed, a total of 4,320 trials per subject per model per comparison (across 15 target images and 8 scaling values) for all energy model comparisons and for luminance model original vs. synthesized white noise comparison. Luminance model synthesized vs. synthesized, white noise comparison had 1,440 trials (across 5 target images and 8 scaling values) for a single subject. Where behavioral data is plotted in this paper, the proportion correct is the average across all relevant trials.

We fit psychophysical curves describing proportion correct as a function of model scaling using the two-parameter function for discriminability *d’* derived in Freeman and Simoncelli (2011):

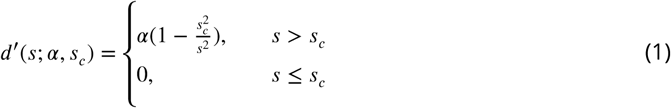

where *s_c_* is the critical scaling value (performance is at chance for scaling values at or below *s_c_*) and *α* is the max *d’* value (called the “proportionality factor” in Freeman and Simoncelli (2011)).

Psychophysical curves were constructed by converting this *d*^f^ into the probability correct using the same function as in Freeman and Simoncelli (2011):

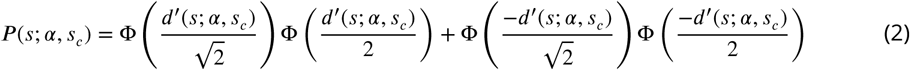

where Ф is the cumulative of the normal distribution. The probability correct is 50% when *d’* = 0 (and thus when scaling is at or below the critical scaling), reaches about 79% when *d’* = 2 and 98% when *d’* = 4. As the α parameter above gives the maximum *d’* value, it has a monotonic relationship with the asymptotic performance, which can be seen in figure 16.

**Figure 16.**
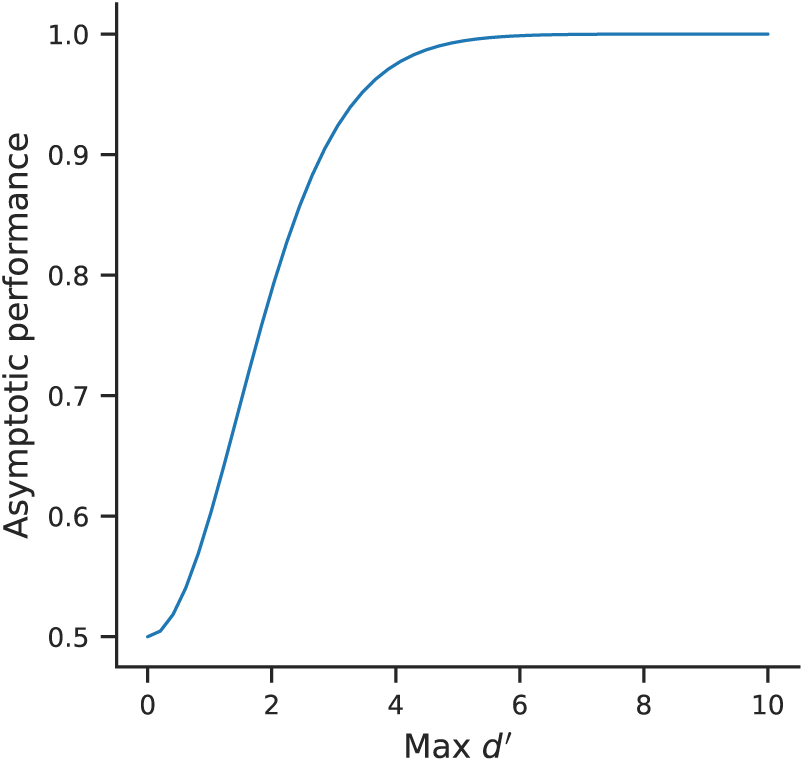
Relationship between the max *d’* parameter, *a* and asymptotic performance. As max *d’* increases beyond approximately 5 (where asymptotic performance is at ceiling), the slope of the psychophysical curve continues to increase (for example, compare the slope of the luminance and energy model original vs. synth white noise comparisons in figure 9A).

The posterior distribution over parameters *s_c_* and α was estimated using a hierarchical, partial-pooling model, with independent subject- and target image-level effects for both *s_c_* and α, with each model and comparison estimated separately, following the procedure used in Wallis et al. (2019). Subject responses were modeled as samples from a Bernoulli distribution with probability (1 – π)*P* (*s*) + 0.5π, where π is the lapse rate, estimated independently for each subject. Estimates were obtained using a Markov Chain Monte Carlo (MCMC) procedure written in Python 3.7.10 (Van Rossum and Drake, 2009) using the numpyro package, version 0.8.0 (Phan et al., 2019; Bingham et al., 2018). MCMC sampling was conducted using the No U-Turn Sampler algorithm (Hoffman and Gelman, 2014), with parameters selected to ensure convergence, which was assessed using the *R^^^* statistic (Brooks and Gelman (1998), looking for *R^^^ <* 1.01, Vehtari et al. (2021)) and by examining traceplots. Parameters were given weakly-informative priors and both *s_c_* and *α* were estimated on natural logarithmic scales.

In sum, for model *m* e {*E, L*}, comparison *t*, subject *x*, target image *i*, and scaling *s*:

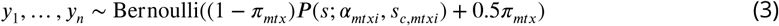

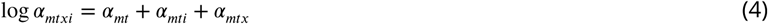

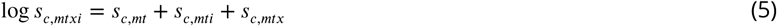

with the following priors:

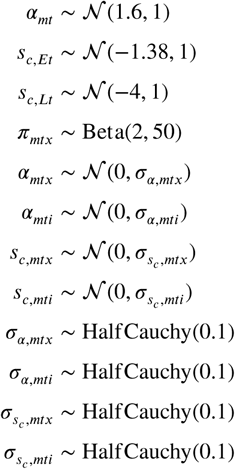

The priors for *s_c,mt_* of the energy and luminance models correspond to critical scales of 0.25 and 0.018, respectively. These values were taken from the centers of the V1 physiological range provided in Freeman and Simoncelli (2011) figure 5 and from the slope of a line fit to the dendritic field diameter vs. eccentricity of midget retinal ganglion cells in Dacey and Petersen (1992) figure 2B. These priors reflect our original prediction that the models’ critical scaling values should be similar to those of the physiological scaling in the brain area sensitive to the same image features. While we are not committed to this link to physiology, it is reasonable to expect the energy model’s critical scaling to be similar to that of Freeman and Simoncelli (2011), who tested the same model, and to assume the critical scaling for the luminance model to be a good deal smaller. The priors also reflect our prediction that critical scaling should be independent of comparison type and consistent across target images and subjects, while not placing too much of a constraint on the parameters (see appendix 7 for a comparison with an unpooled MCMC model, where parameters are estimated independently for each psychophysical curve, and a partial-pooling model that adds interaction terms between subjects and target images).

The posterior distribution represents the model’s beliefs about the parameters given the priors and data and is summarized throughout this paper as the posterior mean and 95% highest density intervals. The latter represents the range of values containing 95% of the distribution with the highest probability, as opposed to the more common 95% confidence interval, which is symmetrically arranged around the mean. The two are identical for symmetric distributions, but can diverge markedly if the distribution is highly skewed (Kruschke, 2015).

### Software

These experiments relied on a variety of custom scripts written in Python 3.7.10 (Van Rossum and Drake, 2009), all found in the GitHub repository associated with this paper. The following packages were used: snakemake (Mölder et al., 2021), JAX (Bradbury et al., 2018), matplotlib (Hunter, 2007), psychopy (Peirce et al., 2019), scipy (Virtanen et al., 2020), scikit-image (van der Walt et al., 2014), pytorch (Paszke et al., 2019), arviz (Kumar et al., 2019), numpyro (Phan et al., 2019; Bingham et al., 2018), pandas (Reback et al., 2021; McKinney, 2010), seaborn (Waskom, 2021), jupyterlab (Kluyver et al., 2016), and xarray (Hoyer and Hamman, 2017).

## Acknowledgments

The authors would like to thank David Brainard for the use of his photographs, from the UPenn Natural Image Database (Tkačik et al., 2011) and the extended dataset of urban images. They would also like to thank Tony Movshon, David Heeger, David Brainard, Corey Ziemba, and Colin Bredenberg for their feedback on the manuscript, Mike Landy for his assistance with the design of the psychophysical task and feedback on the manuscript, Heiko Schütt for his assistance with the Markov Chain Monte Carlo analysis, and the authors of Wallis et al. (2019) for sharing their code and data. Furthermore, they would like to thank Liz Lovero, Paul Murray, Dylan Simon, and Aaron Watters for their work in creating the metamer browser website.

## Footnotes

1 The schematics presented in this and similar figures are meant as conceptual aids to help think through the metamer paradigm and the results presented in this paper. The metameric regions are depicted as discrete tiles, which helps us make several didactic points. A more realistic representation would be low-dimensional surfaces (manifolds) within the high dimensional pixel space.

2 Note that this assumption does not preclude recurrent processing or feedback from later stages or other modalities. It only asserts that these signals do not carry additional information about the current visual stimulus.

