## Appendix for "Foveated metamers of the early visual system"

### Appendix 1

#### Luminance model metamers initialized with natural images

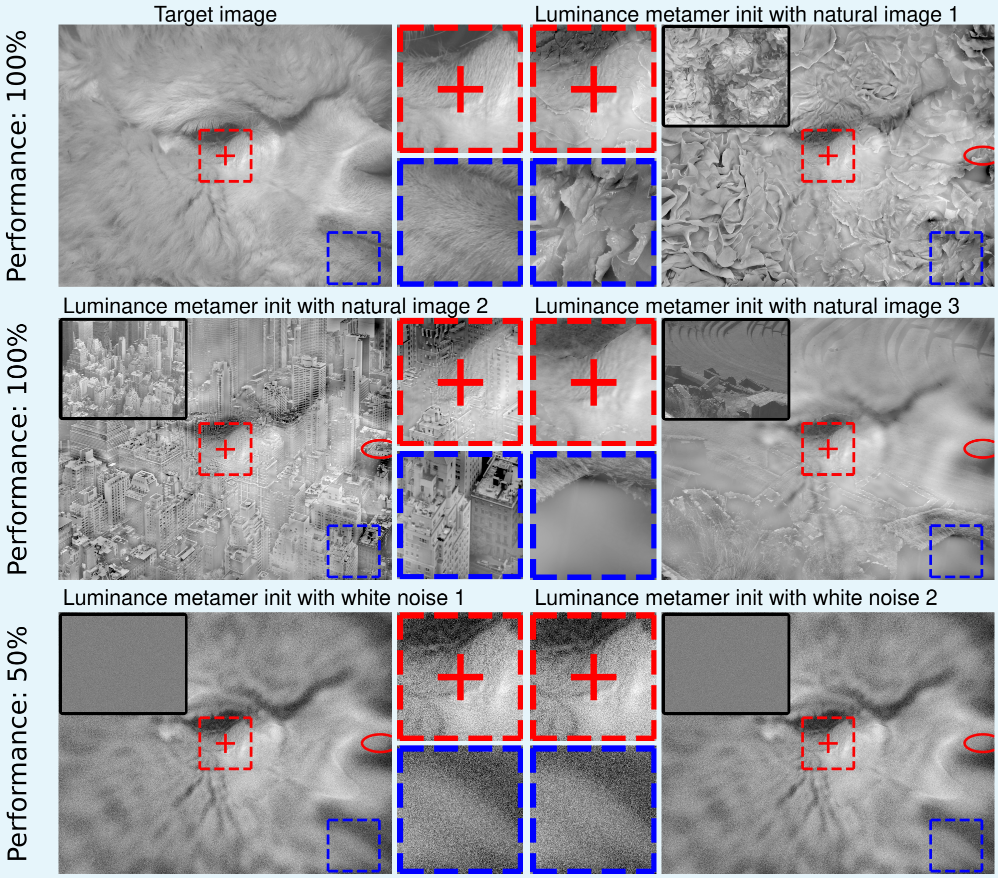


**Appendix 1 Figure 1.** With luminance model metamers synthesized with scaling value 0.23, performance depends strongly on the comparison. We only have data for the comparison in the bottom row (synthesized vs. synthesized white noise), where performance is at chance. From inspection, we believe performance would be at ceiling for either the original vs. synthesized or synthesized vs. synthesized comparisons when initializing with natural images (top right: initialized with lettuce, middle left: nyc, middle right: terrace). Insets show the images used to initialize metamer synthesis. Full resolution version of this figure can be found on the OSF.

Owing to the length of time required to synthesize luminance model metamers with scaling values in the range required for the original vs. synthesized comparison (from 2 days for scaling 0.058 to 14 days for scaling 0.01), we did not generate a full set of luminance model metamers initialized with natural images and thus did not run the corresponding psychophysical experiment. However, we believe we would obtain similar results as those we observed for the energy model: the performance on the original vs. synthesized comparison would be approximately the same, while it would be greatly improved for the synthesized vs. synthesized comparison, with a smaller critical scaling value and a higher max $d'$. Some example comparisons are presented in figure 1 to illustrate this. All presented model metamers were generated with scaling 0.23. The bottom row contains two model metamers initialized with different patches of white noise, while the other three model metamers were initialized with other natural images. We only have data for the comparison in the bottom row, where performance was at chance, but, examining the other stimuli, it is likely that performance would be at ceiling for those comparisons. For all model metamers, the pooling windows have grown to the point that a good deal of information from the initial images remains in the ultimate metamer. If these initial images were patches of white noise, that information is impossible for humans to use for discriminating between the two stimuli, but this is not the case if they were natural images. The luminance model predicts all three comparisons would be equally difficult, but the bottom comparison is impossible, while the other two appear to be trivial (based on examining luminance model metamers initialized with natural images at other scaling values, we believe this is true for the range of scaling values tested for the synthesized vs. synthesized comparison, but would need an experiment to understand scaling values in the range tested for the original vs. synthesized comparison).

### Appendix 2

#### Luminance model metamers initialized with pink noise

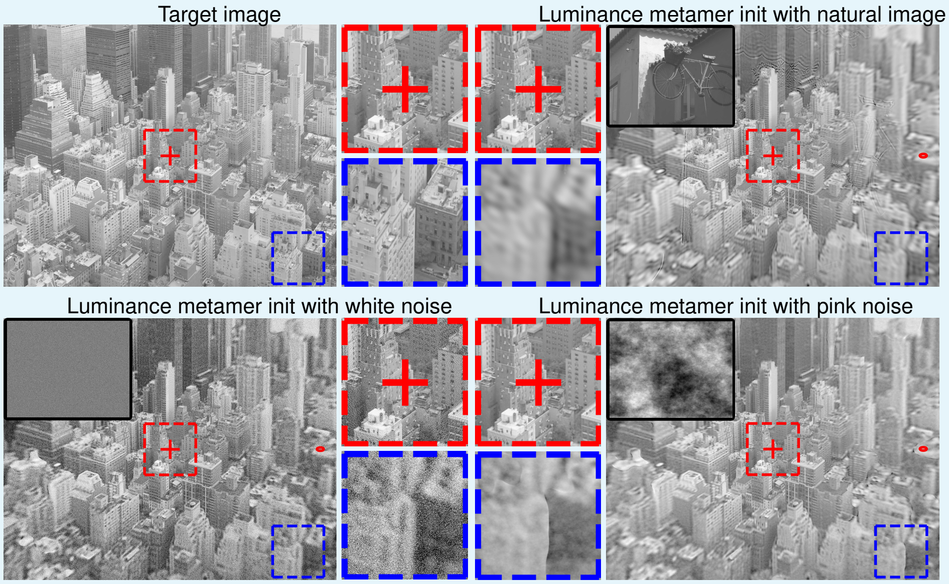


**Appendix 2 Figure 1.** Examples of synthesized luminance model metamers, generated from different initial image types, each with scaling value 0.045. Discriminability from the target image depends on the initialization. For this scaling value, performance was at ceiling when comparing the white noise-initialized model metamer (bottom left) to the target image (top left), as can be seen in figure 6A in the main text. However, the pink noise-initialized metamer (bottom right) lacks the high frequencies used to perform this discrimination; from inspection, we believe performance would be at or near chance. From inspection, we believe performance for the natural image-initialized metamer (top right) would be intermediate between the two. Insets show the images used to initialize metamer synthesis. Full resolution version of this figure can be found on the OSF.

As discussed in the main text, participants make use of the high frequencies present in the luminance model metamers initialized with white noise when distinguishing them from the target images. One might therefore wonder what luminance model metamers initialized with pink noise, which have the same $1/f$ power distribution found in natural images, would look like. As can be seen in figure 2, metamers initialized with pink noise have far less power in the high frequencies compared to those initialized with white noise at the same scaling value. Fixating on the center of these images, the white noise-initialized metamer is more discriminable from the target image than that initialized with pink noise. If a full experiment were run, we believe this comparison would lead to a right-shifted psychophysical curve and a higher critical scaling value. However, the logic of our experiments relies on finding the *smallest* critical scaling value. Thus, while this example shows the importance of synthesis initialization in metamer synthesis, we do not believe it would affect our interpretation of our results or our conclusions.

### Appendix 3

#### Comparison with pooling windows from previous studies

Previous studies computing the critical scaling of pooling models using psychophysical experiments and model metamers have used the raised-cosine windows described in ***Freeman and Simoncelli*** (***2011***) (***Deza et al., 2019***; ***Wallis et al., 2019***). In this study, we use Gaussians instead. Though they match the previously-used windows in their polar arrangement and radial elongation, one might worry that some difference in their construction might influence our results. The Gaussian windows do overlap more substantially than the raised-cosine windows and, being Gaussians, extend infinitely (unlike the raised-cosine windows), leading to a smoother representation, which resulted in higher-quality synthesized stimuli. Use of these models and the scientific inferences made by the authors do not rely on the exact specification of the windows, and we believe our models can be viewed as comparable to previous results (similarly, ***Deza et al.*** (***2019***) construct a texture-like representation using the representation of VGG19, rather than the Portilla-Simoncelli texture statistics (***Portilla and Simoncelli, 2000***) used by ***Freeman and Simoncelli*** (***2011***); ***Wallis et al.*** (***2019***), but their psychophysical results are comparable).

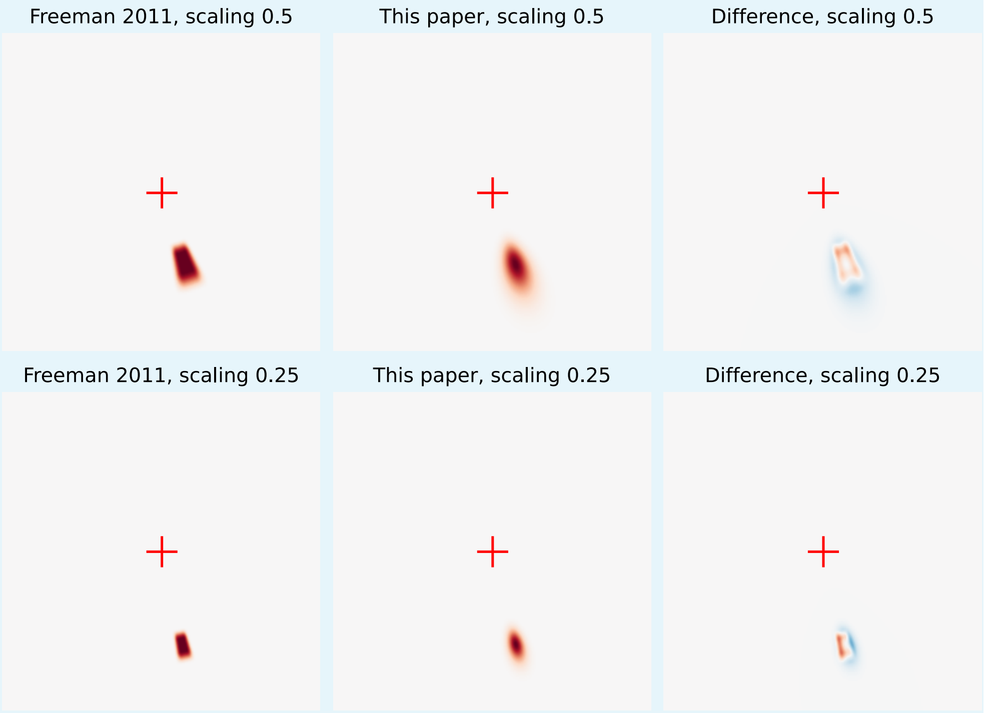


**Appendix 3 Figure 1.** Scaling has the same interpretation for the Gaussian windows of this study and the raised-cosine windows used in previous studies. The left column shows the windows from ***Freeman and Simoncelli*** (***2011***), the center column shows the windows from this paper, and the right column shows their difference. The windows have been normalized so as they both have a maximum value of 1 (when used in our models, the Gaussian windows have a lower maximum value so that the sum across all windows is 1). While the centers and extent of the two sets of windows are different, their full-width at half-maximum is matched.

However, interpretation of these results does critically depend on the definition of the model's scaling parameter, which gives the ratio between the pooling windows' radial full-width at half-maximum and central eccentricity. We preserve that definition, as can be viewed in figure 3, which shows an example window from ***Freeman and Simoncelli*** (***2011***) (constructed using their MATLAB code found on GitHub) and our Gaussian implementation at two scaling values. Both sets of windows were constructed to have a resolution of 512 by 512 pixels, with a maximum eccentricity of 13 and multiple scales; the presented windows are from the second-coarsest scale (thus, a resolution of 256 by 256 pixels) to demonstrate we handled the multi-scale construction similarly as well (similar results are found for other scales).

The left column shows the windows from ***Freeman and Simoncelli*** (***2011***), the center column shows the windows from this paper, and the right column shows their difference. The windows have been normalized so as they both have a maximum value of 1 (when used in our models, the Gaussian windows have a lower maximum value so that the sum across all windows is 1). Several aspects are visible:

- The centers of the windows are not perfectly aligned. The exact sampling of the two sets of windows is slightly different, which should have no impact on their interpretation.
- The visible extent of the Gaussian is slightly larger. As discussed above, Gaussians extend infinitely, whereas the raised-cosine windows have a finite support. This leads to the desired smoother representation.
- However, the full-width at half-maximum (in both directions) appears to be matched. Scaling therefore has the same interpretation for both sets of windows.

### Appendix 4

#### Differences with Freeman and Simoncelli, 2011

As noted in the text, while the critical scaling we found for the energy model comparing two synthesized stimuli initialized with white noise was comparable to that found in ***Freeman and Simoncelli*** (***2011***), subjects' asymptotic performance was lower (average of 60% correct across subjects and target images, compared to 85%). This was also noted, for the "mid-ventral", pooled texture-statistic model, in ***Wallis et al.*** (***2019***), who ran a control experiment to ensure it doesn't come down to task differences. As our implementation of the pooling models is completely separate from that of ***Freeman and Simoncelli*** (***2011***), we investigated whether differences in window construction or other implementation details could have led to any marked difference. A Jupyter notebook investigating this can be found in the Github repo associated with this project, showing that the windows appear comparable, as do energy model metamers with comparable scaling values. Additionally, if our windows were significantly different from those in ***Freeman and Simoncelli*** (***2011***), we would expect them to affect the critical scaling, whose value relates window radial diameter to eccentricity. However, the critical scaling value for the only comparison present in both studies (energy model, synthesized vs. synthesized white noise comparison) is consistent, so we believe it is unlikely that some detail of window construction is responsible for the differences between the studies.

One possible explanation for the difference in asymptotic performance is the smaller pixel pitch of our stimuli: 48.5 pixels per degree, as compared to the 19.7 pixels per degree used in ***Freeman and Simoncelli*** (***2011***). To investigate this possibility, we down-sampled our target images by half (to a resolution of 1024 by 1300 pixels) using a Gaussian pyramid (via scikit-image's `transform.pyramid_reduce` function, ***van der Walt et al.*** (***2014***); ***Burt and Adelson*** (***1987***)) then synthesized energy model metamers with identical scaling values and optimization hyperparameters to the energy model synthesized vs. synthesized white noise comparison. The psychophysical experiment was run as before, with one subject (sub-00), upsampling the stimuli with nearest neighbor interpolation to present at the same physical size as before, effectively doubling the pixel pitch.

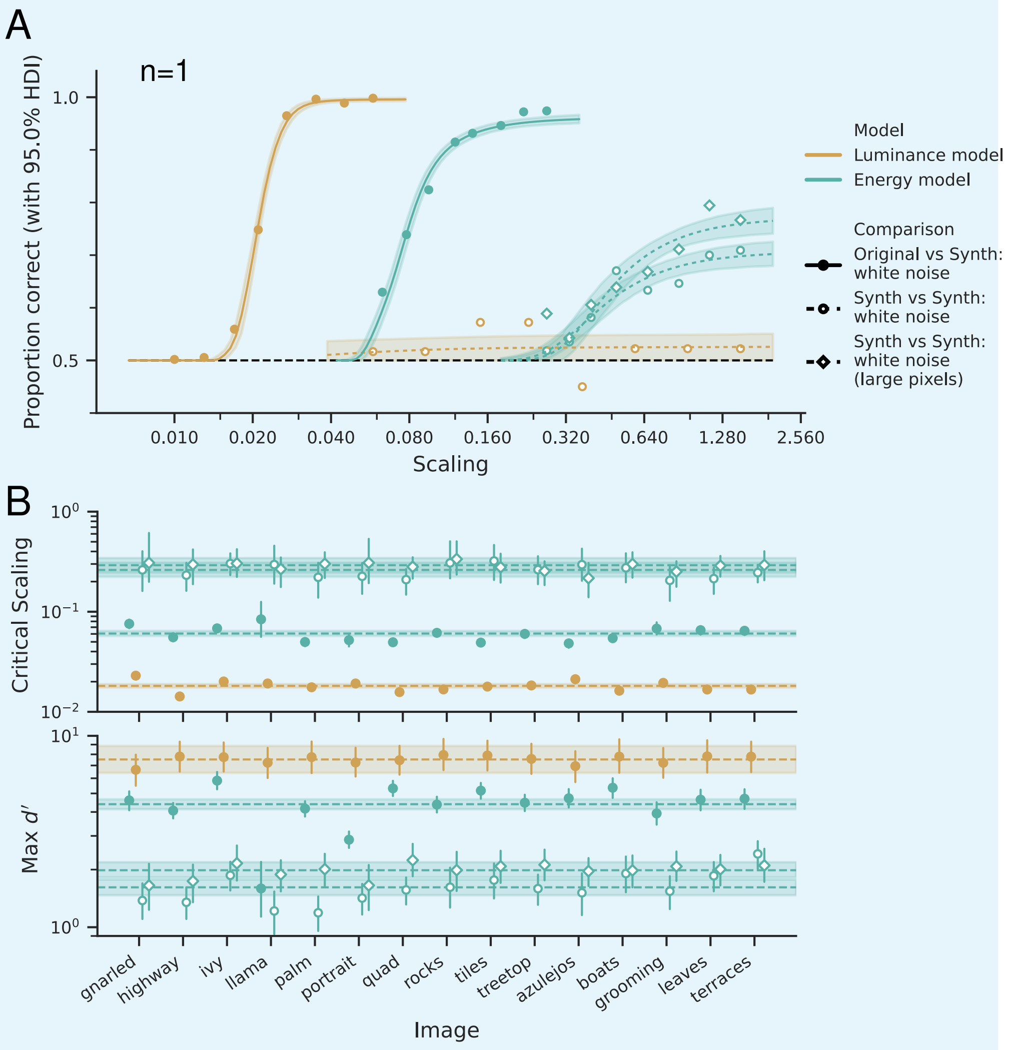


**Appendix 4 Figure 1.** Asymptotic performance on the synthesized vs. synthesized white noise comparison is slightly higher when stimuli have a larger pixel pitch. **(A)** Probability correct for one subject, sub-00, as a function of scaling for energy and luminance models. Data points represent the average across target images, 540 trials per data point (one subject, fifteen target images) except for luminance model synthesized vs. synthesized: white noise comparison, which have 180 per data point (one subject, five target images). Lines represent the posterior predictive means across target images, with the shaded region giving the 95% HDI. **(B)** Parameter values for these comparisons. Top row shows the critical scaling value and the bottom the value of the max $d'$ parameter. Left column presents the values for each target image separately for this one subject, while the right presents the values for this subject, averaged across target images. In this case, we only present the data for sub-00, as they are the only subject to perform the task with larger pixel pitch. Points represent the posterior means, shaded regions the 95% HDI, and horizontal dashed lines and shaded regions average across all shown target images for this subject. Note that the luminance model, synthesized vs. synthesized: white noise comparison is not shown in this panel, because the data was poorly fit by this curve — as can be seen in panel A, the psychophysical curve is essentially flat at chance and thus the fit had low max $d'$ and low critical scaling), with high uncertainty.

As can be seen in appendix 4 figure 4, which shows data for a single subject, the critical scaling value did not change, but asymptotic performance increased slightly. For an intuition, see the example stimuli in appendix 4 figure 1, which show that the patterns used to differentiate the two synthesized stimuli are slightly more obvious with a larger pixel pitch.

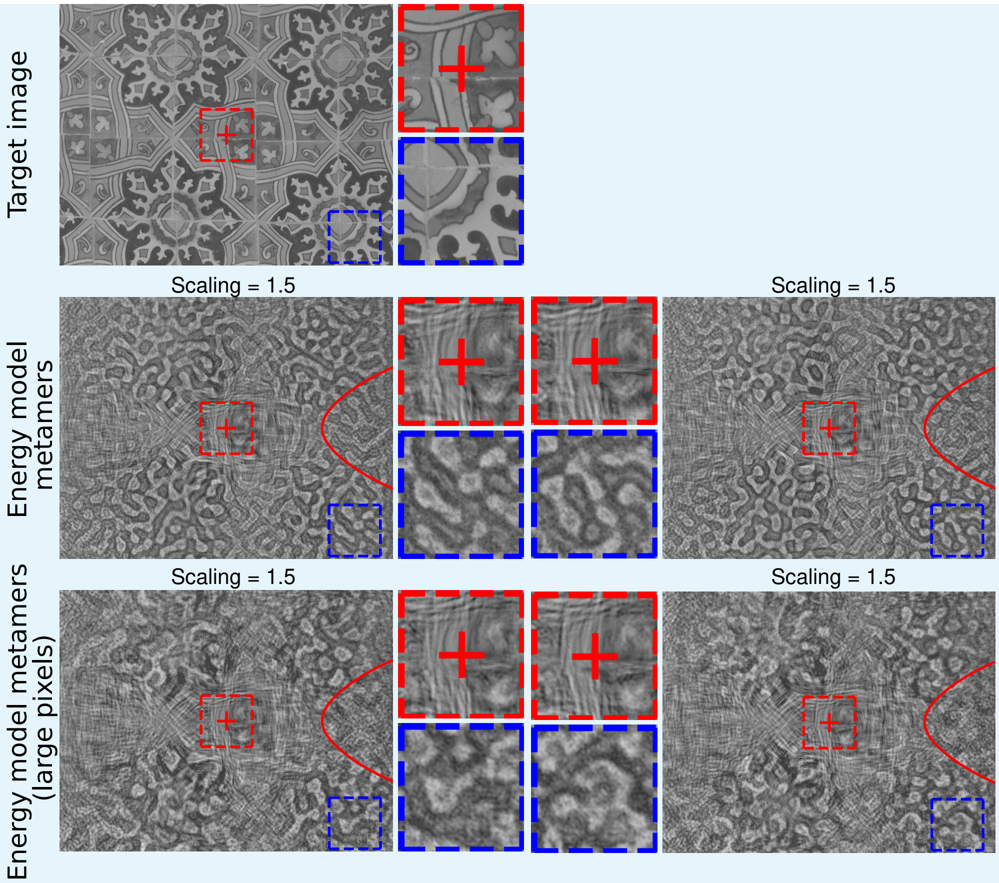


**Appendix 4 Figure 2.** Energy model metamers synthesized with larger pixel pitch are slightly easier to distinguish from each other at high scaling values. Top row shows the target image stimulus whose representation all presented model metamers were synthesized to match. Middle row shows two energy model metamers initialized with different patches of white noise, which were shown in the synthesized vs. synthesized white noise comparison. Bottom row shows two model metamers initialized with same settings but half the pixel resolution, which were shown in the large pixel version of the synthesized vs. synthesized white noise comparison. As these stimuli have half the resolution in each direction as the middle row but were presented at the same physical display size, their pixel pitch was doubled, making it comparable to that of ***Freeman and Simoncelli*** (***2011***). With the larger pixel pitch, the snake-like patterns that can be used to distinguish the two synthesized stimuli have a lower spatial frequency (in cycles per degree), which, given their presence in the periphery where spatial frequency sensitivities are lower, may account for the participant's increased ability to distinguish such stimuli. Full resolution version of this figure can be found on the OSF.

Beyond the difference in pixel pitch, there are several other potential factors that may contribute to our lower asymptotic performance:

- Experimental parameters were determined for the original vs. synthesized comparison, which is an easier task than synthesized vs. synthesized comparison. Longer presentation times or some other configuration may increase asymptotic performance in synthesized vs. synthesized comparison (but, as ***Freeman and Simoncelli*** (***2011***) showed, these sorts of experimental manipulations are unlikely to have much effect on the critical scaling, which is the focus of this study).
- Our stimuli are physically larger, with a diameter of 53.6 degrees compared to 26 degrees in ***Freeman and Simoncelli*** (***2011***). In debriefing, participants reported performing the task by finding particular informative parts of the stimulus (e.g., high contrast edges) and attending there. Larger stimuli may have made these regions harder to find, and the attentional manipulation in ***Freeman and Simoncelli*** (***2011***) shows that

attending to the most informative region of the stimulus improves asymptotic performance (while leaving critical scaling unchanged). Additionally, ***Ziemba and Simoncelli*** (***2021***) showed that the probability of correctly differentiating two samples from the same texture family decreases as those samples get larger (conversely, performance increases with stimulus size when participants are differentiating between two different texture families). Something analogous may be happening here, with two synthesized stimuli initialized with white noise acting similarly to two samples from the same texture family.

- As seen both here and in previous studies (***Freeman and Simoncelli, 2011***; ***Wallis et al., 2019***; ***Deza et al., 2019***), asymptotic performance varies more across target images and subjects than critical scaling. We have a different subset of target images and subjects and so sampling issues may be partly at fault.

All told, there are multiple reasons why asymptotic performance differs between this study and ***Freeman and Simoncelli*** (***2011***), which we are unable to comprehensively track down.

### Appendix 5

#### Model metamers are physically distinct from their target stimulus

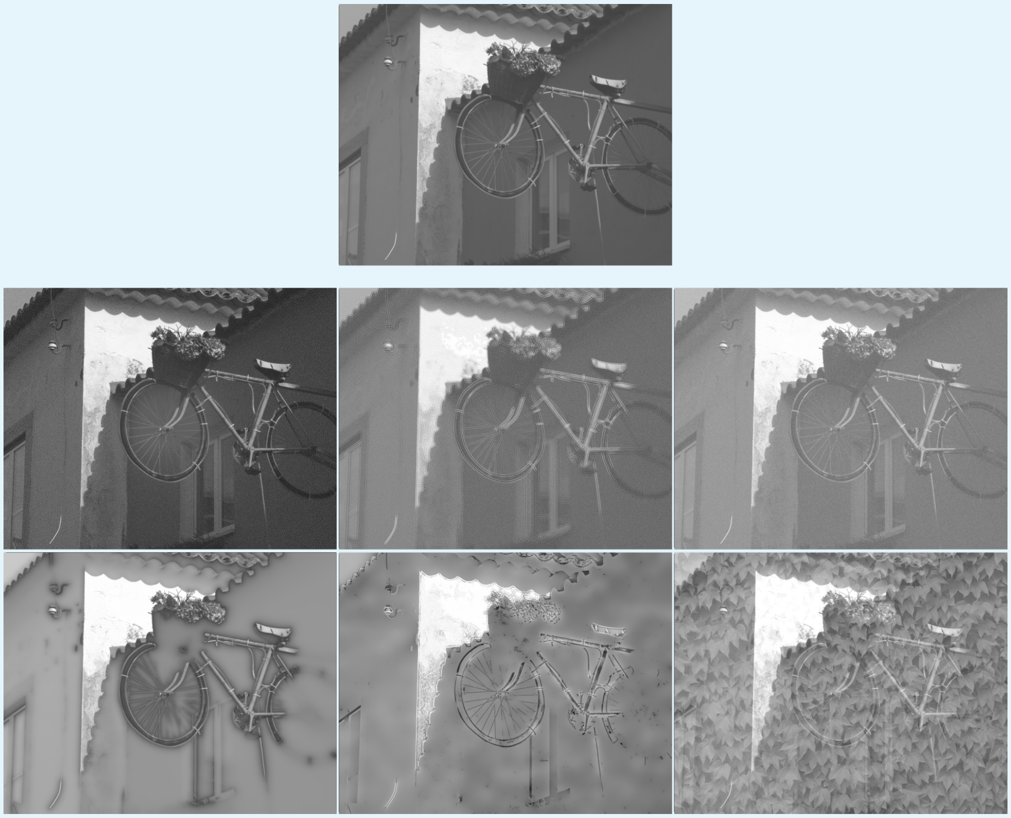

**Appendix 5 Figure 1.** Luminance model metamers are substantially physically different from their target images. The six bottom stimuli all have approximately the same mean-squared error to the top image, the target natural image. Notice that the luminance model metamer synthesized with scaling value 0.01 (top left) is by far the least discriminable with the natural image when fixating on the center, the others are obviously distinct. This pair of original and synthesized image was chosen to set the MSE because it has the lowest MSE for all luminance metamers when compared with their target image. Full resolution version of this figure can be found on the OSF.

There are two properties that perceptual metamers must have: they must be perceptually identical and physically distinct. We have shown our synthesized stimuli are perceptually identical and figure 5 shows that the stimuli are physically distinct. The image at the top of that figure is the original natural image whose representation the model metamers match. The six lower images all have approximately the same mean-squared error to that original image, yet only the luminance model metamer, top left, is confusable with the natural image; the others are easily discriminable from it. The top right image has had high-frequency noise added uniformly across the image and thus shows the importance of foveation: the luminance model metamer also differs from the original image primarily by the addition of high-frequency noise but, by concentrating the noise in the periphery, it is undetectable.

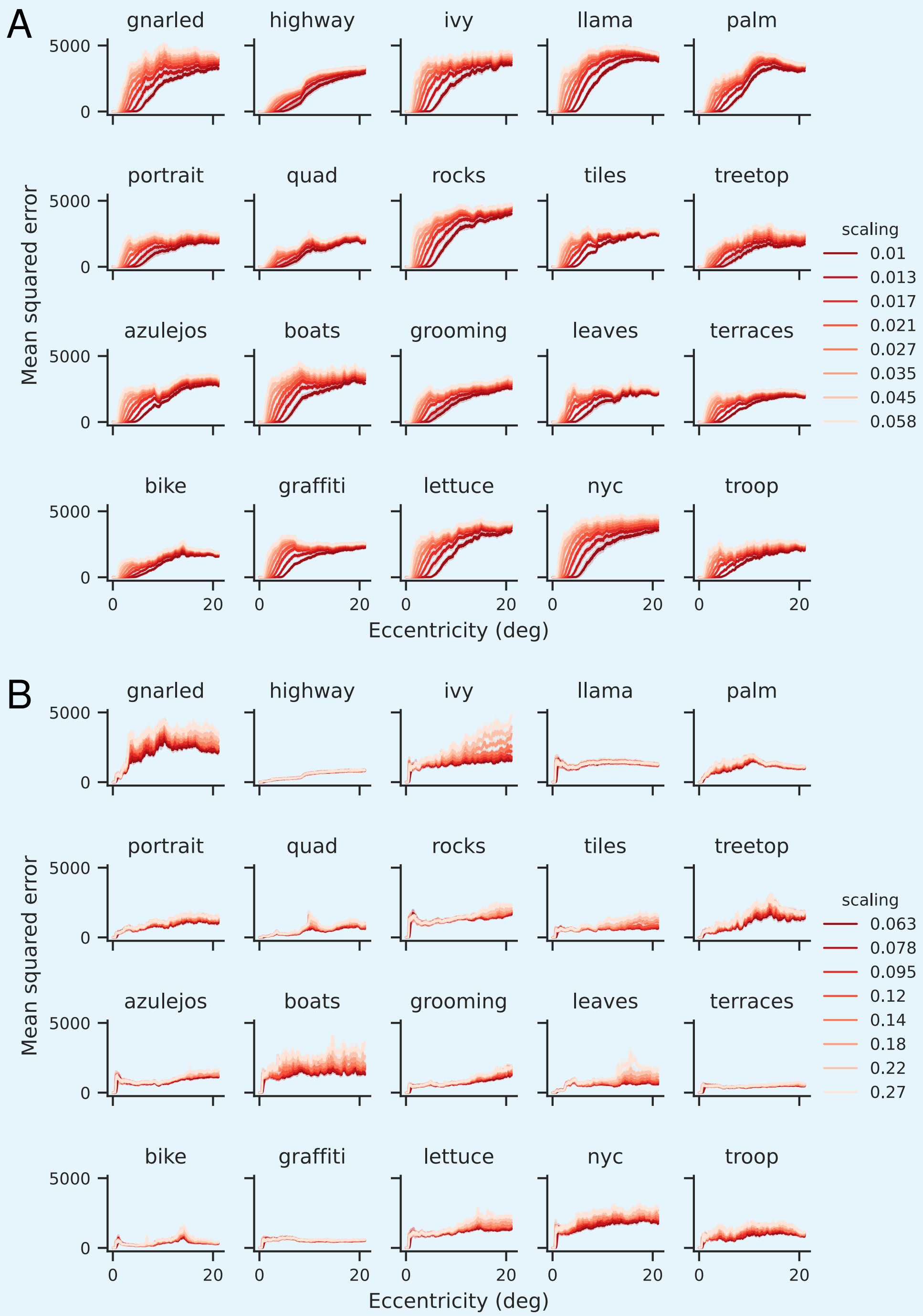


**Appendix 5 Figure 2.** Mean-squared error (MSE) betwen model metamers and target images as a function of eccentricity, averaged radially, for the luminance model **(A)** and energy model **(B)**. MSE is computed between images represented as 8-bit unsigned integers. Each scaling value is shown as a separate color, and each target image is plotted on a separate axis. For the luminance model, the smallest scaling value has zero MSE out to about 5 degrees, and all scaling values rise with eccentricity and then saturate. For the energy model, all scaling values have non-zero MSE by 1 degree and are indistinguishable beyond that for most images, though for some, such as gnarled and ivy, lower scaling values have higher MSE across eccentricities.

In addition to global mean-squared error, we can also examine the mean-squared error at each eccentricity, as plotted in figure 1, which shows the mean-squared error between model metamers and target images as a function of eccentricity for the range of scaling values used in the original vs. synthesized comparison, for the luminance model (A) and the energy model (B). As discussed earlier, when windows get smaller than a pixel, the only value that will have the same model output is the original pixel value. One might have a similar concern for the energy model: what is the smallest pooling window in which a given spatial frequency can be computed and so will windows containing e.g., 16 pixels, also be uniquely constrained when matching 6 scales? Panel (B) shows this is not the case for our energy model metamers: all scaling values have a non-zero mean-squared error by 1 degree. However, panel (A) shows that the lowest scaling value for the luminance model metamers has a zero mean-squared error until about 5 degrees, and rises beyond that. While lower scaling values do have a smaller mean-squared error across much of the image, that does not guarantee that they are less informative: mean-squared error is a poor perceptual metric (***Wang and Bovik, 2009***).

However, when mean-squared error is zero, two stimuli cannot be discriminated between, and so we might worry that the reduced performance for lower scaling values of the luminance model does not reflect the fact that we have found perceptual metamers but that we have removed all information that participants could use to discriminate between the stimuli because of sampling issues with our display. We think this is unlikely for two reasons. First, participants are able to use peripheral information to discriminate between stimuli, including letters (***Song et al., 2014***), gratings, and Vernier lines (***Duncan and Boynton, 2003***). Second, our resolution is approximately 48.5 pixels per degree, giving a Nyquist frequency of about 24 cycles per degree. Human grating acuity drops below this frequency by an eccentricity of 2 or 3 degrees (***Duncan and Boynton, 2003***; ***Anderson et al., 1991***), suggesting that participants would not be able to use this information even if it were present. Additionally, participants reported that the most informative portions of the stimulus were in the mid-periphery, across all conditions, providing additional evidence that they were not relying on the portion of the stimuli where the lowest scaling values were matched to the target stimulus in order to perform the task. Further experiments investigating which portions of the stimulus are the most informative in a more systematic way, such as restricting the stimuli to annuli at different eccentricities and comparing the resulting psychophysical curves, would provide clarity on this matter.

### Appendix 6

#### Target image and subject differences

Figures 6 and 1 show the performance for each target image and subject separately, respectively (collapsing over the other dimension). As can also be seen in figure 10, there's not much variability in subjects' critical scaling.

We also see that the two extreme target images (examined in figure 8) are only extremes for the energy model when comparing against natural and synthesized stimuli: with the exception of the top right panel, which is replotted from 8, llama and nyc no longer appear as the two ends of the continuum, but lie instead in the middle. In the synthesized vs. synthesized comparisons, the elongated hard edges are not present in either stimulus. For the luminance model original vs. synthesized comparison, elongated contours are no harder to capture than any other natural image feature. In either case, the nyc target is no longer special. Likewise, the llama is not an especially hard comparison for the luminance model, because when the pooling windows get large, the synthesized stimulus contains mostly white noise whereas the original image stimulus contains mostly cloud-like patterns.

We can also see from this figure that the between-image differences are largest for the original vs. synthetic energy model, compared to the other three. For the original vs. synthesized luminance model comparison, this is because there is less interaction between the original image features and the model's invariances: the model is insensitive to high frequencies and thus requires smaller windows to adequately represent that information, but all of our target images are natural images, with $1/f$ power distributions, and thus there is relatively little information present in higher frequencies. For the energy model synthesized vs. synthesized comparison, the two stimuli to distinguish are both synthesized and thus neither contain the edge information that allows subjects to readily distinguish the original from synthesized stimulus in the original vs. synthesized comparison.

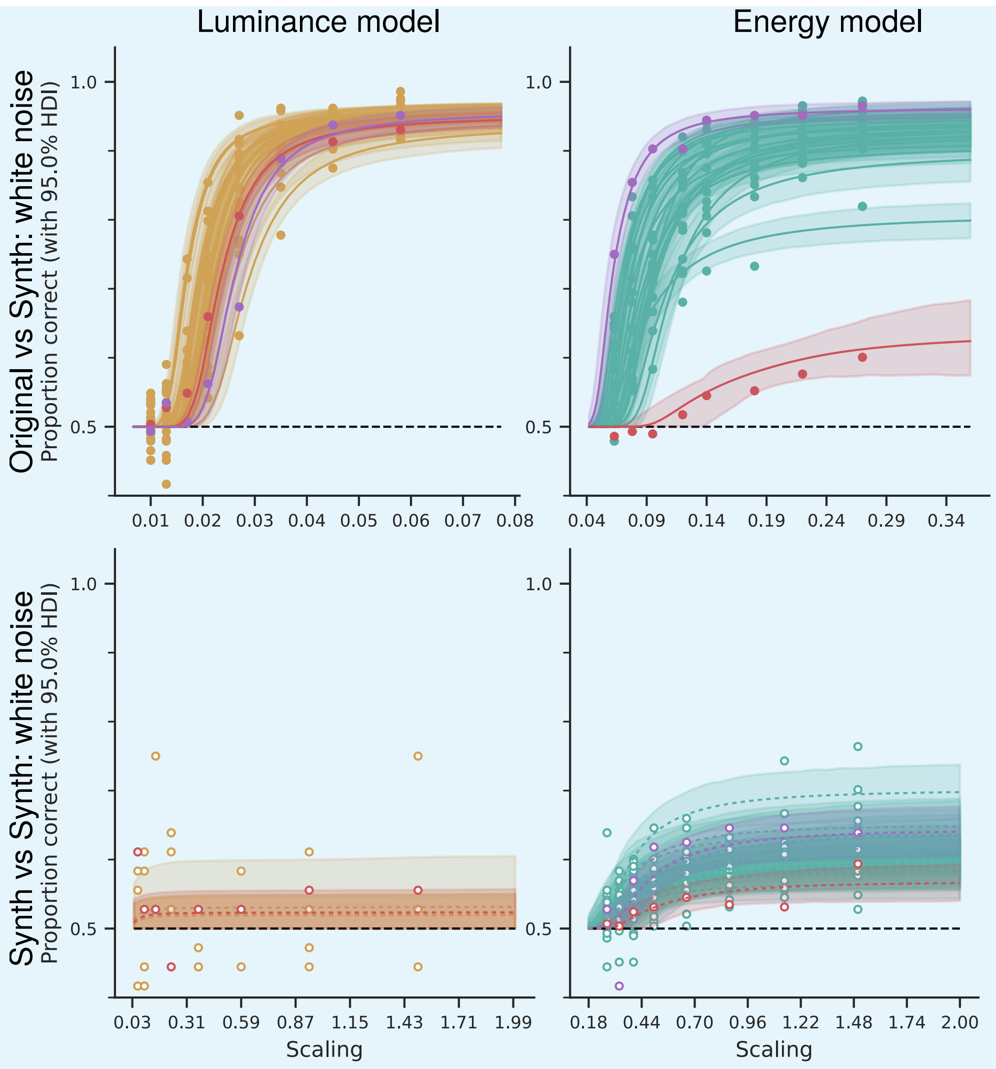


**Appendix 6 Figure 1.** Image content only matters for the energy model, original vs. synthesized comparison. Performance for each target image, averaged across subjects, is plotted separately. This is because the image content affects performance based on how it interacts with the models' sensitivities and insensitivities: llama contains little phase structure and is thus easy for the energy model to capture even at larger scaling values, while nyc contains many hard edges, and is thus difficult for the energy model to capture even with the smallest tested windows. However, for the luminance model original vs. synthesized comparison, the relevant interaction would be with the Fourier spectra: the model is insensitive to high frequencies in its periphery. As all images have $1/f$ spectra everywhere in the image, none are outliers in terms of difficulty. For the synthesized vs. synthesized comparisons, on the other hand, by definition, neither displayed stimulus contains those features that the model is insensitive to, which is what enables correct performance in the original vs. synthesized case. Thus, stimuli are more interchangeable with each other. Colors and symbols are the same as in 6, except for the two lines showing the nyc (purple) and llama (red) target images, which are colored as in 8. Top right subplot is reprinted from 8A. Note that bottom left subplot only contains five target images (and in particular does not include the purple line representing nyc), as only a single session of that comparison was run.

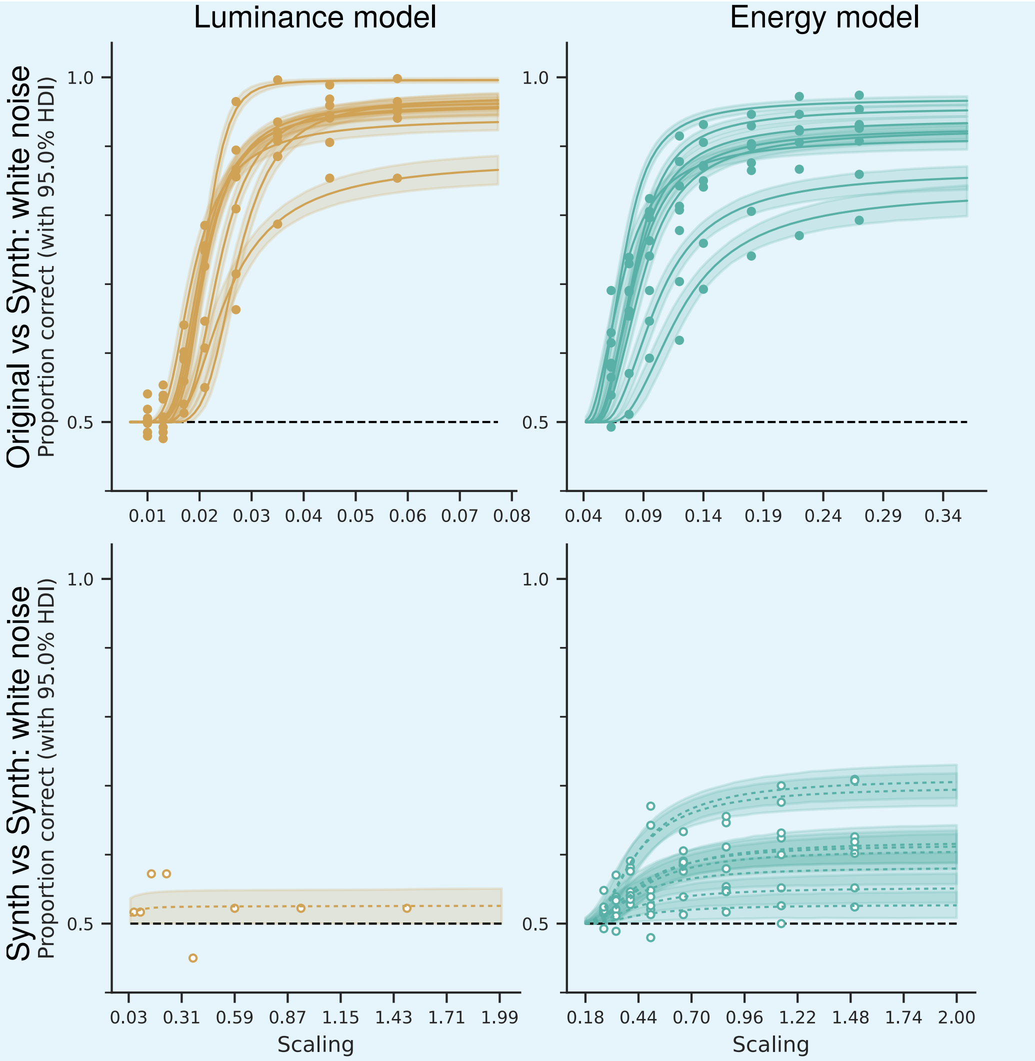


**Appendix 6 Figure 2.** Performance does not vary much across subjects, and varies more in terms of max $d'$ than critical scaling. Each line represents the performance for a single subject, averaged across target images. Colors and symbols are the same as in 6. Note that bottom left subplot only contains a single subject, since only one subject completed this comparison.

### Appendix 7

#### MCMC model comparison

The model used to fit our psychophysical curves assumes that each parameter (max $d'$, $\alpha$, and critical scaling, $s_c$) has a global mean along with subject- and target image-level effects, with no interaction between the two. While this parameterization keeps the total number of parameters very low (29 for each of max $d'$ and critical scaling: one for the global mean, 20 for the image-level, and 8 for the subject-level) and allows for a subject to be particularly good at the task (resulting in a higher max $d'$ or a lower critical scaling) or a target image to be particularly difficult, it does not allow for an individual subject to find a target image particularly difficult. To investigate whether this was biasing our results, we fit two additional parameterizations: a completely unpooled model, where max $d'$ and critical scaling were estimated independently for each psychophysical curve (i.e., each model, comparison, subject, and target image) and a partially-pooled model with interactions, which takes the model used in the main paper, as described in the methods section (see equation 3) and adds interaction terms.

That is, for model $m \in \{E, L\}$, comparison $t$, subject $x$, target image $i$, and scaling $s$, the unpooled model fits performance using the parameterization:

$$y_1, \ldots, y_n \sim \text{Bernoulli}((1 - \pi_{mtx})P(s; \alpha_{mtxi}, s_{c,mtxi}) + 0.5\pi_{mtx}) \tag{6}$$

with priors:

$$\begin{aligned} \alpha_{mtxi} &\sim \mathcal{N}(1.6, 1) \\ s_{c,Etxi} &\sim \mathcal{N}(-1.38, 1) \\ s_{c,Ltxi} &\sim \mathcal{N}(-4, 1) \\ \pi_{mtx} &\sim \text{Beta}(2, 50) \end{aligned}$$

While the interactions model uses the parameterization:

$$y_1, \ldots, y_n \sim \text{Bernoulli}((1 - \pi_{mtx})P(s; \alpha_{mtxi}, s_{c,mtxi}) + 0.5\pi_{mtx}) \tag{7}$$

$$\log \alpha_{mtxi} = \alpha_{mt} + \alpha_{mti} + \alpha_{mtx} + \alpha_{mtxi} \tag{8}$$

$$\log s_{c,mtxi} = s_{c,mt} + s_{c,mti} + s_{c,mtx} + s_{c,mtxi} \tag{9}$$

with the following priors:

$$
\begin{aligned}
\alpha_{mt} &\sim \mathcal{N}(1.6, 1)\\
s_{c,Et} &\sim \mathcal{N}(-1.38, 1)\\
s_{c,Lt} &\sim \mathcal{N}(-4, 1)\\
\pi_{mtx} &\sim \mathrm{Beta}(2, 50)\\
\alpha_{mtx} &\sim \mathcal{N}(0, \sigma_{\alpha,mtx})\\
\alpha_{mti} &\sim \mathcal{N}(0, \sigma_{\alpha,mti})\\
\alpha_{mtxi} &\sim \mathcal{N}(0, \sigma_{\alpha,mtxi})\\
s_{c,mtx} &\sim \mathcal{N}(0, \sigma_{s_c,mtx})\\
s_{c,mti} &\sim \mathcal{N}(0, \sigma_{s_c,mti})\\
s_{c,mtxi} &\sim \mathcal{N}(0, \sigma_{s_c,mtxi})\\
\sigma_{\alpha,mtx} &\sim \mathrm{HalfCauchy}(.1)\\
\sigma_{\alpha,mti} &\sim \mathrm{HalfCauchy}(.1)\\
\sigma_{\alpha,mtxi} &\sim \mathrm{HalfCauchy}(.1)\\
\sigma_{s_c,mtx} &\sim \mathrm{HalfCauchy}(.1)\\
\sigma_{s_c,mti} &\sim \mathrm{HalfCauchy}(.1)\\
\sigma_{s_c,mtxi} &\sim \mathrm{HalfCauchy}(.1)
\end{aligned}
$$

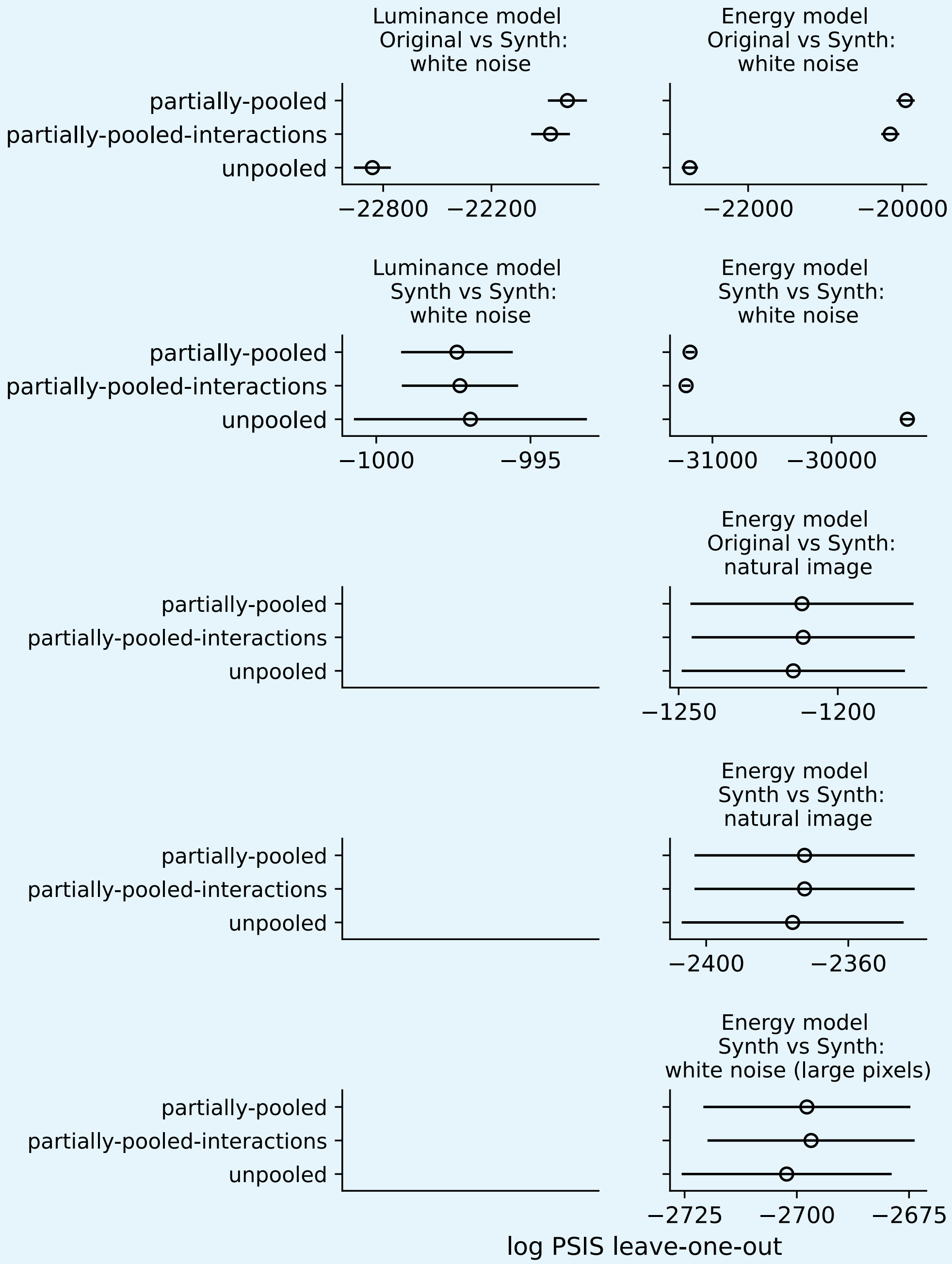


**Appendix 7 Figure 1.** Comparison of models used to fit psychophysical curves, using log-scored leave-one-out cross-validation, estimated using Pareto-smoothed importance sampling (***Vehtari et al., 2016***). A higher score corresponds to better predictive accuracy. Each model was fit separately to each model and comparison. The models differ in how parameters are modeled across subjects and target images: the "unpooled" model assumes no relationship across subjects or target images, the "partially-pooled" model has separate image- and subject-level effects for each parameter, and the "partially-pooled-interactions" model adds interaction terms to that model.

A comparison between the three models for all comparisons for which data was collected can be seen in figure 7, using log-scored leave-one-out cross-validation, estimated using Pareto-smoothed importance sampling (***Vehtari et al.*** (***2016***), comparable results were obtained using the widely applicable criterion, WAIC, ***Watanabe*** (***2013***)). For the four comparisons for which only a single subject's data was collected (luminance model: synth vs. synth white noise, energy model: original vs. synth natural image, synth vs. synth natural image, and synth vs. synth white noise (large pixels)), the three models perform comparably. With only a single subject, the models do not differ enough to distinguish their predictions from each other.

Of the remaining three comparisons, the partially-pooled model performs best for the two original vs. synth white noise comparisons, while the unpooled model predicts best for the energy model synth vs. synth white noise comparison. Somewhat surprisingly, the interactions model performs slightly worse than the partially-pooled model for each comparison, including the one where they are both outperformed by the unpooled model. This may be because the interactions model has far more parameters to fit than the partially-pooled model, and suggests that the interaction terms are generally unnecessary.

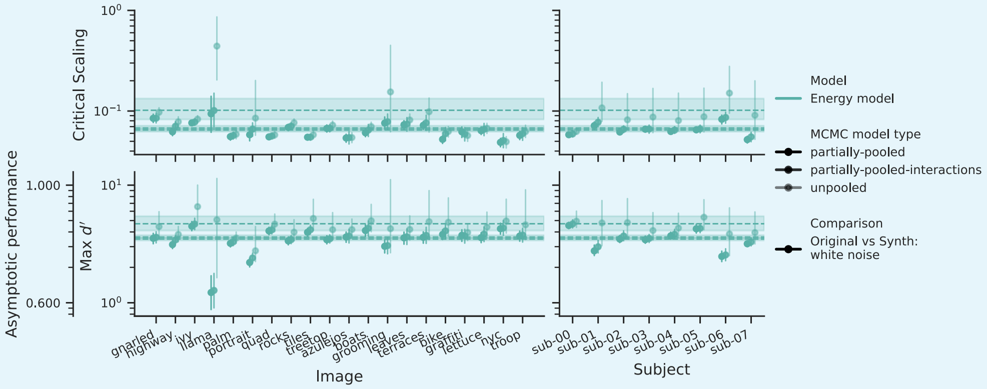


**Appendix 7 Figure 2.** Comparison of parameters for the energy model, original vs. synth white noise comparison, averaged across subjects and target images, for each of the three MCMC models. The partially-pooled values are identical to those shown in figure 6. The values for the unpooled model are always greater than or equal than those for the interactions model, which are almost identical to those for the partially-pooled model (though they are, as a rule, slightly larger).

Across all target images and subjects, the unpooled model's parameter estimates are higher than those of the interactions model, followed by the partially-pooled model, though the last two are almost identical (see figure 1). We will thus discuss the unpooled and partially-pooled models. For this comparison, the critical scaling values differ most for the llama, grooming, and terraces images; the others are quite similar. Individual performance curves for both MCMC models are plotted in figure 2, which may elucidate these differences.

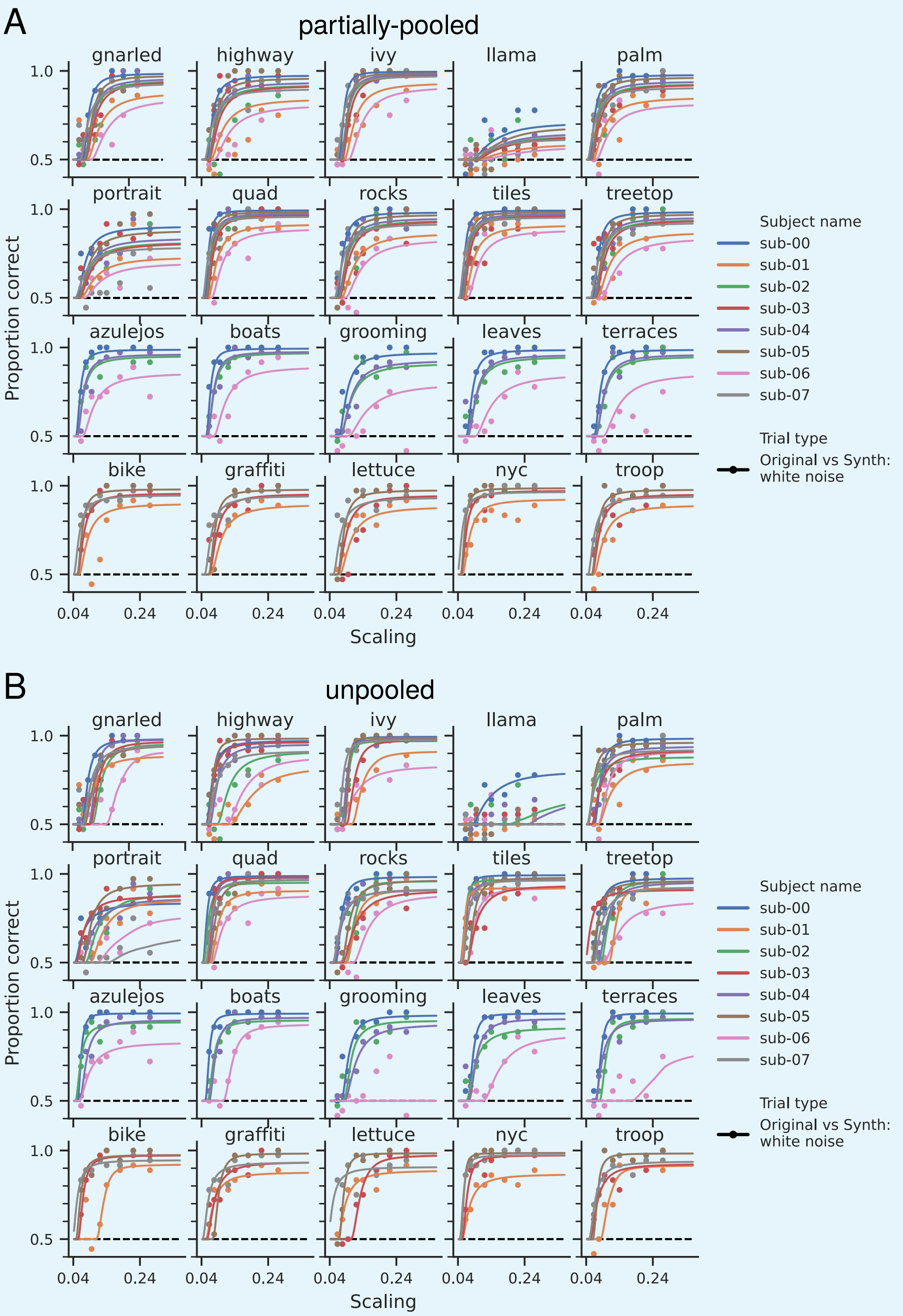


**Appendix 7 Figure 3.** Comparison between psychophysical curves fit by the partially-pooled and unpooled MCMC models for the energy model original vs. synth comparison. The two models diverge the most when performance data appears ambiguous and, in general, the unpooled model fits higher values for both the max $d'$ and critical scaling.

The difference between these two MCMC models arises when there is ambiguous data: when performance data could be fit using either a lower critical scaling and max $d'$ or a higher critical scaling and max $d'$ value, the unpooled model picks the latter, the partially-pooled the former. These two parameter regimes make different predictions about subject performance as the synthesis model's scaling grows beyond the tested values: the unpooled model predicts that performance will continue to increase, while the partially-pooled predicts that performance has stabilized. This can be seen most strikingly in the predictions for the llama image: as discussed earlier, this target image is an outlier in its difficulty for all subjects. The two models differ in how they interpret this difficulty: the unpooled model predicts that asymptotic performance is comparable to those of other target images, once we synthesize model metamers with higher scaling values, while the partially-pooled model predicts performance will not increase much beyond the observed values. Based on the reasons outlined earlier in this paper, we believe the partially-pooled predictions are the most reasonable. This can also be seen in the diverging predictions for sub-06's performance on the grooming image, where they performed below chance for the highest tested scaling value. The partially-pooled model essentially ignores this data point, assuming it's largely due to lapses, while the unpooled model interprets this performance as requiring a significantly higher critical scaling value. Again, the predictions of the partially-pooled model seem more reasonable.

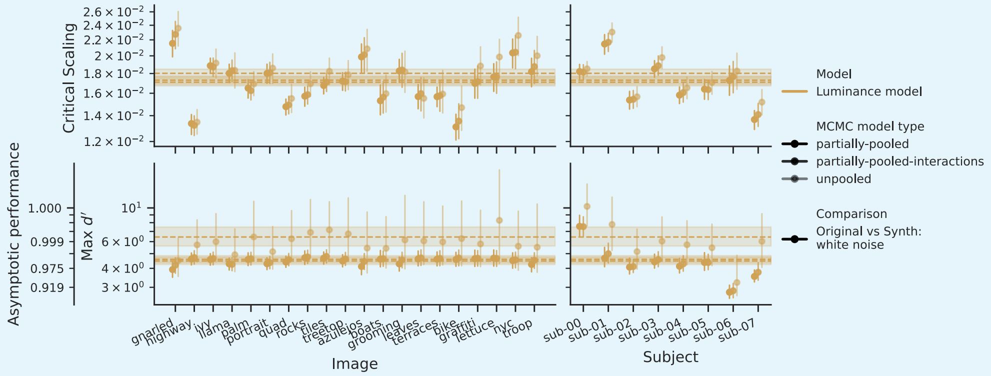


**Appendix 7 Figure 4.** Comparison of parameters for the luminance model, original vs. synth white noise comparison, averaged across subjects and target images, for each of the three MCMC models. The partially-pooled values are identical to those shown in figure 6. The values for the unpooled model are always greater than or equal than those for the interactions model, which are almost identical to those for the partially-pooled model (though they are, as a rule, slightly larger).

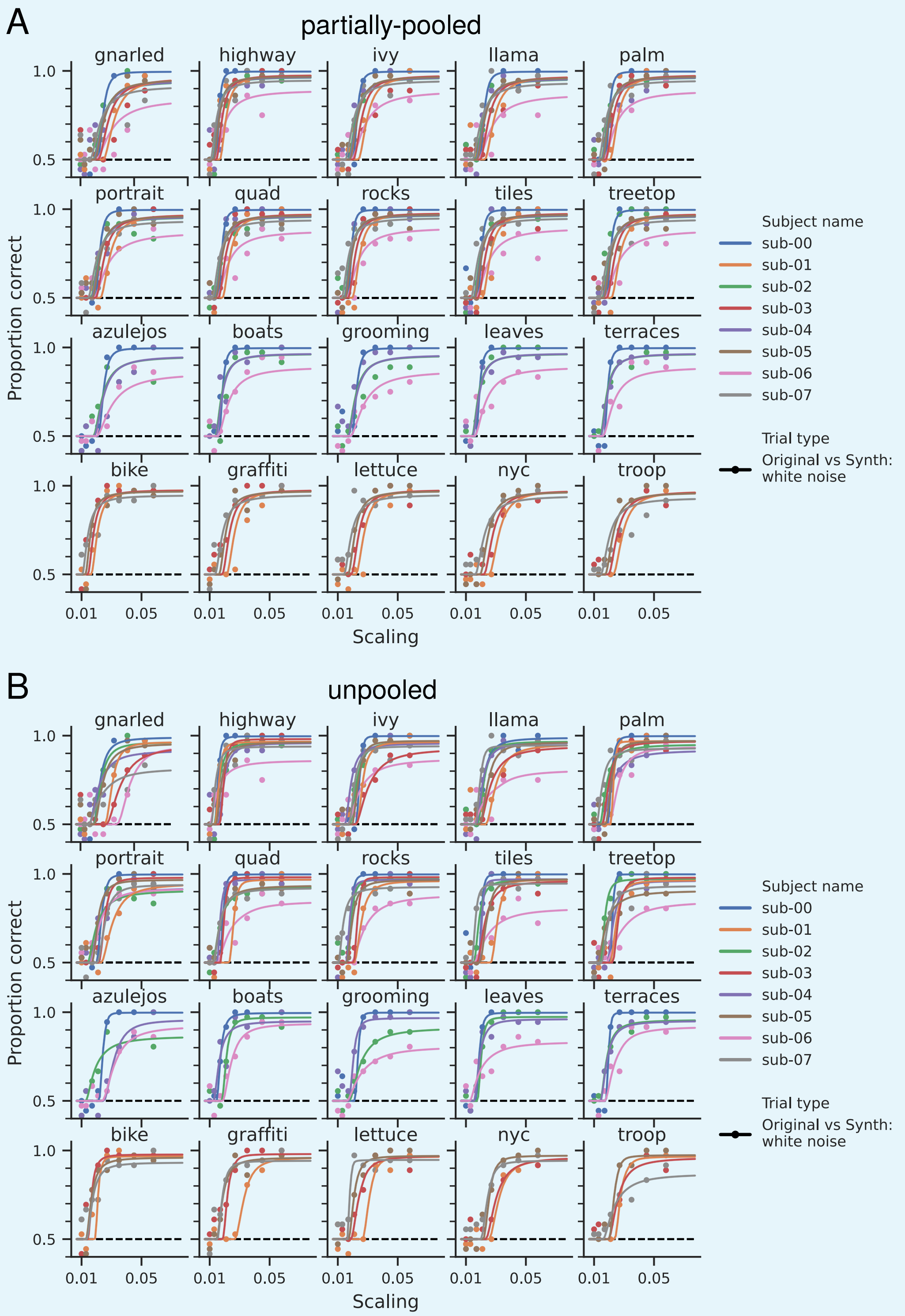


**Appendix 7 Figure 5.** Comparison between psychophysical curves fit by the partially-pooled and unpooled MCMC models for the luminance model original vs. synth comparison. The two models diverge the most when performance data appears ambiguous and, in general, the unpooled model fits higher values for both the max $d'$ and critical scaling.

The same pattern can be seen in the other two comparisons. For the luminance model original vs. synth comparison (parameters in figure 3 and performance in figure 4), the differences in critical scaling are smaller across MCMC models than in the same comparison for the energy model. However, the same basic pattern appears: unpooled parameter estimates are larger than those of the interactions model, which are larger than those for the partially-pooled model.

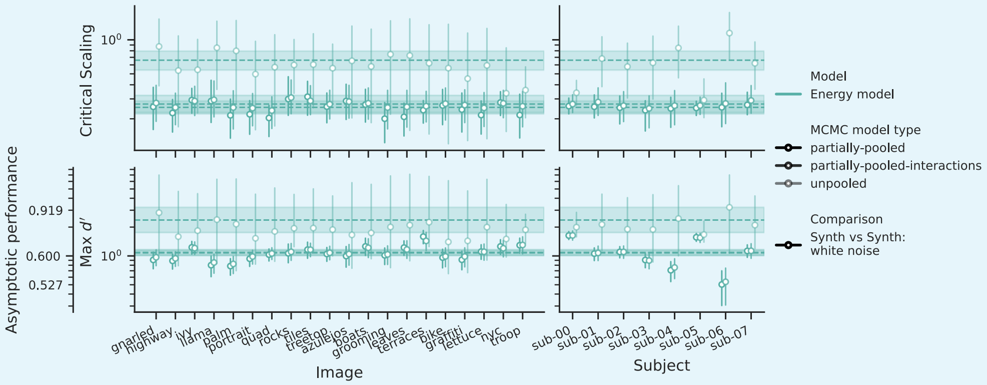


**Appendix 7 Figure 6.** Comparison of parameters for the energy model, synth vs. synth white noise comparison, averaged across subjects and target images, for each of the three MCMC models. The partially-pooled values are identical to those shown in figure 6. The values for the unpooled model are always greater than or equal than those for the interactions model, which are almost identical to those for the partially-pooled model (though they are, as a rule, slightly larger).

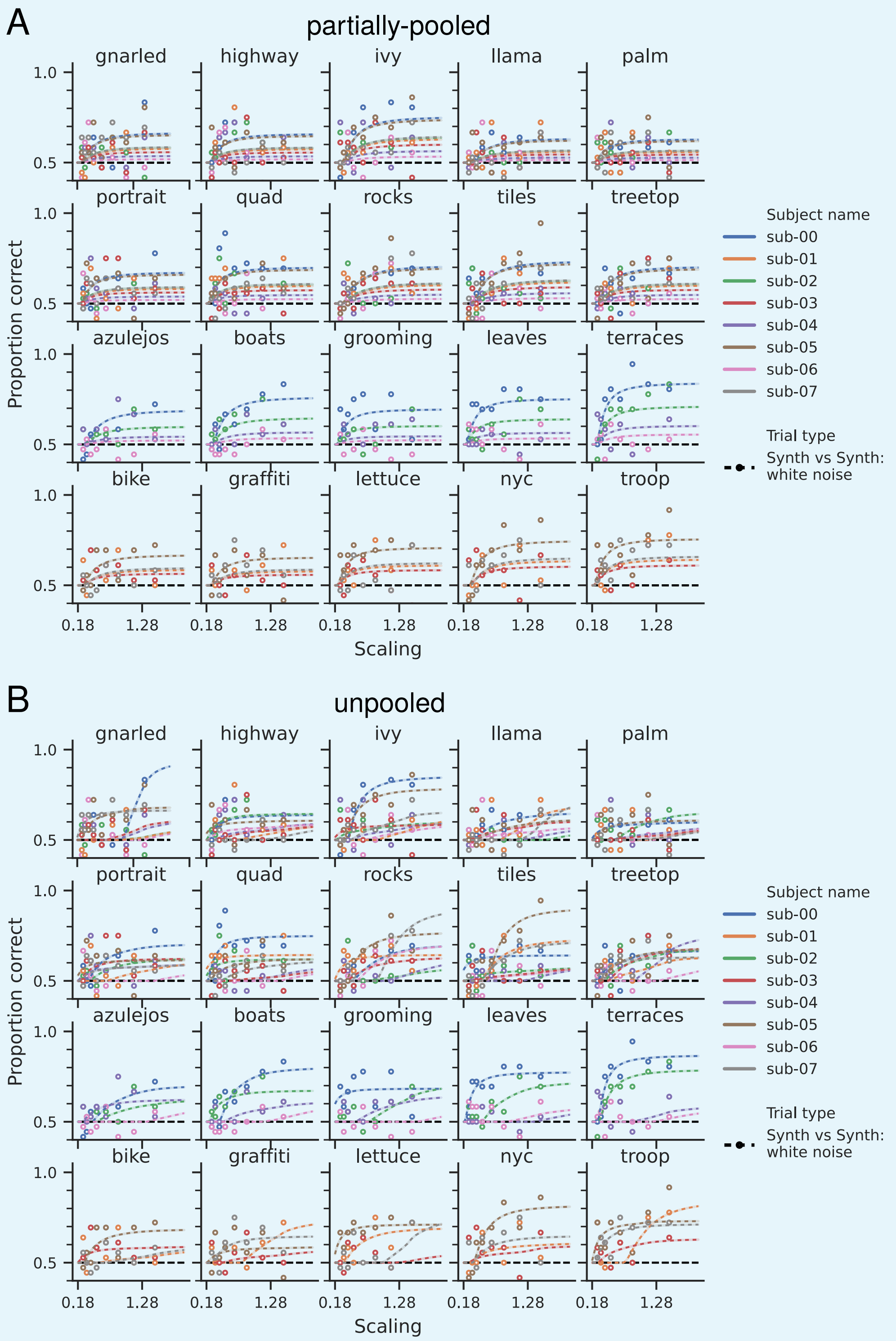


**Appendix 7 Figure 7.** Comparison between psychophysical curves fit by the partially-pooled and unpooled MCMC models for the energy model synth vs. synth comparison. The two models diverge the most when performance data appears ambiguous and, in general, the unpooled model fits higher values for both the max $d'$ and critical scaling.

The largest difference across MCMC models is found in the energy model synth vs. synth comparison (parameters in figure 5 and performance in figure 6), which is also the only comparison in which the unpooled model performs better than the partially-pooled one.

As can be seen by looking at the data in figure 6, the data quality is poorer than that of the original vs. synth comparisons, which explains the larger discrepancy between the MCMC models' fitted parameters. However, the unpooled model fit a comparable max $d'$ to the other comparisons (and thus a higher critical scaling), whereas the partially-pooled model fits both parameters lower. As discussed in section Critical scaling is smaller for original vs. synthesized comparisons than synthesized vs. synthesized comparisons, we believe that asymptotic performance is indeed lower for this comparison and that further increasing the synthesizing model's scaling will not lead to performances approaching 90% accuracy, as the unpooled model predicts. We thus believe that the partially-pooled model's parameters are more accurate, despite the unpooled model's better cross-validation performance.

We thus believe that our use of the partially-pooled model, and its parameters, is justified in this paper. Furthermore, we should note that none of our conclusions change substantially if using a different MCMC model: in all cases, the luminance model original vs. synth comparison has the smallest critical scaling, followed by the energy model original vs. synth and the energy model synth vs. synth; initializing with a natural image still has an effect on the synth vs. synth critical scaling, but not that of the original vs. synth value. A larger critical scaling value for the energy model synth vs. synth does mean that our value would no longer align with that of ***Freeman and Simoncelli*** (***2011***), but that is the only change.

Additionally, this model comparison exercise highlights an important point: while our conclusions do not depend on the subjects' asymptotic performance (as discussed in Why does this comparison-type dependency decrease with feature complexity?, the metamer paradigm makes no predictions for discriminable stimuli), a higher asymptotic performance does make the fitting of the psychophysical curves easier and thus reduces the uncertainty on the inferred parameter values (as we saw that the greatest divergence in parameter values across models comes from curves whose asymptotic performance was poor). As described in appendix 4, there are many reasons why our asymptotic performance may be lower in this energy model synth vs. synth comparison, some of which may be due to the experimental paradigm. In this paper, we used consistent experimental protocols across all comparisons, but the importance of asymptotic performance to obtaining good quality fits suggests that perhaps future studies should consider varying experimental protocols depending on task difficulty in order to obtain comparable uncertainty in parameter values across comparisons, rather than identical experimental paradigms. Care would need to be taken, however, to ensure that the paradigm does not change so much as to affect the interpretation of the models (such as by e.g., removing the blank screen between stimulus presentations or allowing free movement of the eyes).
